# Endothelial AGO1 Drives Vascular Inflammation and Atherosclerosis via a Non-Canonical Nuclear Mechanism

**DOI:** 10.1101/2025.05.01.651783

**Authors:** Xuejing Liu, Dongqiang Yuan, Yingjun Luo, Xiaofang Tang, Alonso Tapia, Naseeb Kaur Malhi, Ting-Yun Wang, Sachchidanand Tiwari, Rahuljeet Singh Chadha, Piotr Swiderski, Marcin Kortylewski, Norbert Pardi, Lu Wei, Satdarshan Pal Singh Monga, Wendong Huang, Kuei-Chun Wang, Zhen Bouman Chen

## Abstract

**BACKGROUND:** Endothelial cell (EC) dysfunction is a cause and consequence of vascular inflammation and lipid dysregulation in atherosclerosis, yet the molecular drivers linking EC dysfunction to systemic metabolic derangements remain incompletely understood. Moreover, whether inhibiting an endogenous gene in ECs can impact liver function, lipid profile, and the vascular inflammation in the context of atherosclerosis has not been demonstrated. We previously identified Argonaute 1 (AGO1), a component of the RNA-induced silencing complex, as a regulator of EC function in angiogenesis and obesity. However, the role of endothelial AGO1 in vascular inflammation and liver function in the context of hyperlipidemia and atherosclerosis is unknown.

**METHODS:** EC-conditional AGO1 knockout (EC-AGO1-KO) and wildtype mice were subjected to pro-atherosclerotic models induced by AAV9-PCSK9 coupled with a Western diet or partial carotid ligation. Metabolic and vascular phenotype and gene expression were analyzed. In human liver sinusoidal and aortic ECs, AGO1 was knocked down using antisense oligonucleotides (ASO), followed by assessment of inflammatory responses (qPCR, RNA-seq, ELISA, and monocyte adhesion assays). To identify the molecular mechanisms linking AGO1 and EC inflammation, Cut&Tag sequencing, chromatin immunoprecipitation, immunofluorescence, proximal ligation assay, and co-immunoprecipitation were performed. The therapeutic effect of AGO1 inhibition was assessed using ASO-delivered via lipid nanoparticle (LNP) for systemic distribution and monocyte membrane-coated nanoparticles (MoNP) to target the inflamed endothelium.

**RESULTS:** EC-AGO1-KO mice exhibited improved plasma lipid profiles, reduced hepatic steatosis, inflammation, and fibrosis, and decreased aortic atherosclerotic burden. AGO1 knockdown in ECs attenuated inflammatory responses. Mechanistically, AGO1 interacted with NF-κB p65 and promoted p65 nuclear translocation and the transcriptional activation of pro-inflammatory genes, including *ICAM1* and *THBS1*. AGO1-ASO delivered through LNP or MoNP achieved the anti-inflammatory, anti-hyperlipidemic, and anti-atherosclerotic effects, recapitulating the phenotypes observed with EC-AGO1-KO.

**CONCLUSIONS:** Endothelial AGO1 promotes vascular inflammation and liver dysfunction in the context of hyperlipidemia and atherosclerosis, in part through a non-canonical nuclear action of AGO1 as an NF-κB coactivator. Inhibition of endothelial AGO1 provides the dual benefits of ameliorating lipid dysregulation and suppressing vascular inflammation. These results highlight EC-AGO1 as a possible therapeutic target for atherosclerosis and cardiometabolic diseases.

## INTRODUCTION

Atherosclerosis, a major cause of cardiovascular disease, arises from hypercholesterolemia due to liver dysfunction and from chronic vascular inflammation.^1^ One of the earliest events that contributes to atherosclerosis is the dysfunction of endothelial cells (ECs), which form the inner lining of the blood vascular wall.^2^ In the classic model of atherosclerosis, cholesterol-loaded low-density lipoproteins (LDLs) become oxidized in the arterial intima and increase expression of adhesion molecules such as intercellular adhesion molecule 1 (ICAM1) and vascular cell adhesion molecule 1 (VCAM1). The upregulation of these adhesion molecules promotes the recruitment and residence of leukocytes, e.g., monocytes, which further differentiate into macrophages, propagating the pro-inflammatory cascade.^3^

The crucial role of EC dysfunction in atherosclerosis is well recognized and substantiated by a large body of literature. Various mouse models with EC-conditional deletion or overexpression of genes encoding adhesion molecules, membrane receptors, adaptor proteins, and transcription factors (TFs) have been characterized for vascular inflammation and remodeling and altered atherosclerotic burden.^3^ The underlying mechanisms include, but are not limited to, the regulation of oxidative stress,^4,5^ nitric oxide (NO) bioavailability,^6^ leukocyte adhesion,^7^ endothelial permeability,^8,9^ and endothelial-to-mesenchymal transition.^10^ Moreover, targeting EC dysfunction using pharmacological approaches such as nanoparticles or EC-directed peptides can attenuate the atherosclerotic lesions in mice.^11–14^ These findings underscore the critical role of endothelial function in atherosclerosis and suggest that targeting ECs may provide therapies to treat atherosclerosis.

Few studies have addressed the role of EC dysfunction in the development of hyperlipidemia in the context of atherosclerosis. Mice with EC-specific gene perturbation typically develop exacerbated or attenuated atherosclerotic lesions without changes in their lipid profile. Given their extensive surface area and angiocrine, cytokine, and metabolic crosstalk with hepatocytes in the highly vascularized liver,^15–18^ EC dysfunction may contribute to dysregulated liver function in atherosclerosis. In support of this view, EC-specific loss of VCAM1 attenuates hepatic inflammation and fibrosis, although without changing lipid metabolism in mice fed a metabolic dysfunction-associated steatohepatitis (MASH) diet.^19^ Deletion or knockdown of EC glucagon-like peptide 1 receptor (*Glp1r)* in mice with MASH abrogated semaglutide’s hepatic benefits independent from weight loss.^20^ Overexpression of human ATP binding cassette subfamily A member 1 (ABCA1), a key membrane transporter for cholesterol efflux in ECs, elevates plasma high-density lipoprotein (HDL) and mitigates atherosclerosis induced by a high-fat, high-cholesterol diet in mice.^21^ However, whether inhibiting an endogenous gene in ECs can impact liver function, lipid profile, and the vascular inflammation in the context of atherosclerosis has not been demonstrated.

In mammalian cells, Argonaute 1 (AGO1) has been characterized mostly as a key protein of the RNA-induced silencing complex (RISC),^22^ with a less characterized nuclear function to regulate gene transcription.^23^ Unlike its homologous AGO2 protein, AGO1 does not possess endonuclease activity to cleave mRNAs. While global AGO2-knockout (KO) mice display severe developmental abnormalities and are embryonically lethal,^24^ AGO1-KO mice are viable.^25^ We found that hypoxia downregulates AGO1 in ECs to promote angiogenesis, in part through microRNA (miR)-mediated post-transcriptional regulation of pro-angiogenic vascular endothelial growth factor A (VEGFA) and angiostatic and pro-inflammatory thrombospondin (TSP1, encoded by *THBS1*).^26,27^ Furthermore, mice with EC-AGO1 deficiency (EC-AGO1-KO) develop an anti-obesity phenotype with increased fat browning and insulin sensitivity.^27^ These findings and recent reports showing an active role of ECs in modulating metabolic function,^19,28–30^ support the idea that ameliorating EC dysfunction may improve metabolic state.^28,31^ Thus, we investigated the effects of inhibition of EC-AGO1 on liver dysfunction and vascular inflammation, as well as the underlying molecular and cellular mechanisms.

In this study, EC-AGO1-KO mice were subjected to an atherogenic regimen and their lipid profiles, liver function, and atherosclerotic lesions were characterized. The effect of AGO1 inhibition in human liver sinusoidal ECs (HLSEC) and human aortic ECs (HAEC), two EC types affected in hypercholesterolemia and atherosclerosis, was determined. EC-AGO1 inhibition in the liver and in the aortic wall *in vivo* and *in vitro* resulted in an anti-atherosclerotic phenotype. Mechanistically, we identified a non-canonical nuclear role of AGO1, by interacting with NF-κB to induce the expression of pro-inflammatory genes in ECs. Systemic inhibition of AGO1 by using lipid nanoparticle (LNP)-encapsulated AGO1-ASO improved hyperlipidemia, liver function, and atherosclerosis. Furthermore, inhibition of EC-AGO1 via a monocyte membrane-coated nanoparticle (MoNP) targeting inflamed endothelium recapitulated the therapeutic effects observed with LNP. Collectively, our findings highlight EC-AGO1 as a critical regulator in atherosclerosis through metabolic and vascular modulations. Targeting EC-AGO1 may confer both metabolic and vascular benefits.

## RESULTS

### EC-AGO1-KO mice show improved lipid profile and liver function under metabolic stress

We previously established a mouse line by crossing *Cdh5-cre* and *AGO1-floxed* mice to study the effect of EC-AGO1-KO on obesity and adipose tissue function.^27^ When subjected to an obesogenic high-fat, high-sucrose (HFHS) diet, along with the previously observed anti-obesity phenotype,^27^ EC-AGO1-KO mice exhibited lower levels of total cholesterol (TC) and low-density lipoprotein/very-low-density lipoprotein (LDL/VLDL) cholesterol, and higher levels of HDL cholesterol, with comparable triglyceride (TG) levels (**Figure S1**). Given the focus on atherosclerosis in this study, we validated that AGO1 was reduced in ECs from the liver and aorta, but not in bone marrow-derived monocytes or peripheral blood mononuclear cells (PBMC) from the EC-AGO1-KO mice compared to wildtype (WT) littermates (**Figure 1A, Figure S2A-H**). Of note, the expression of *Ago2*, another AGO family member more intensively studied, was not affected by AGO1-KO in ECs (**Figure S2I**).

**Figure 1.**
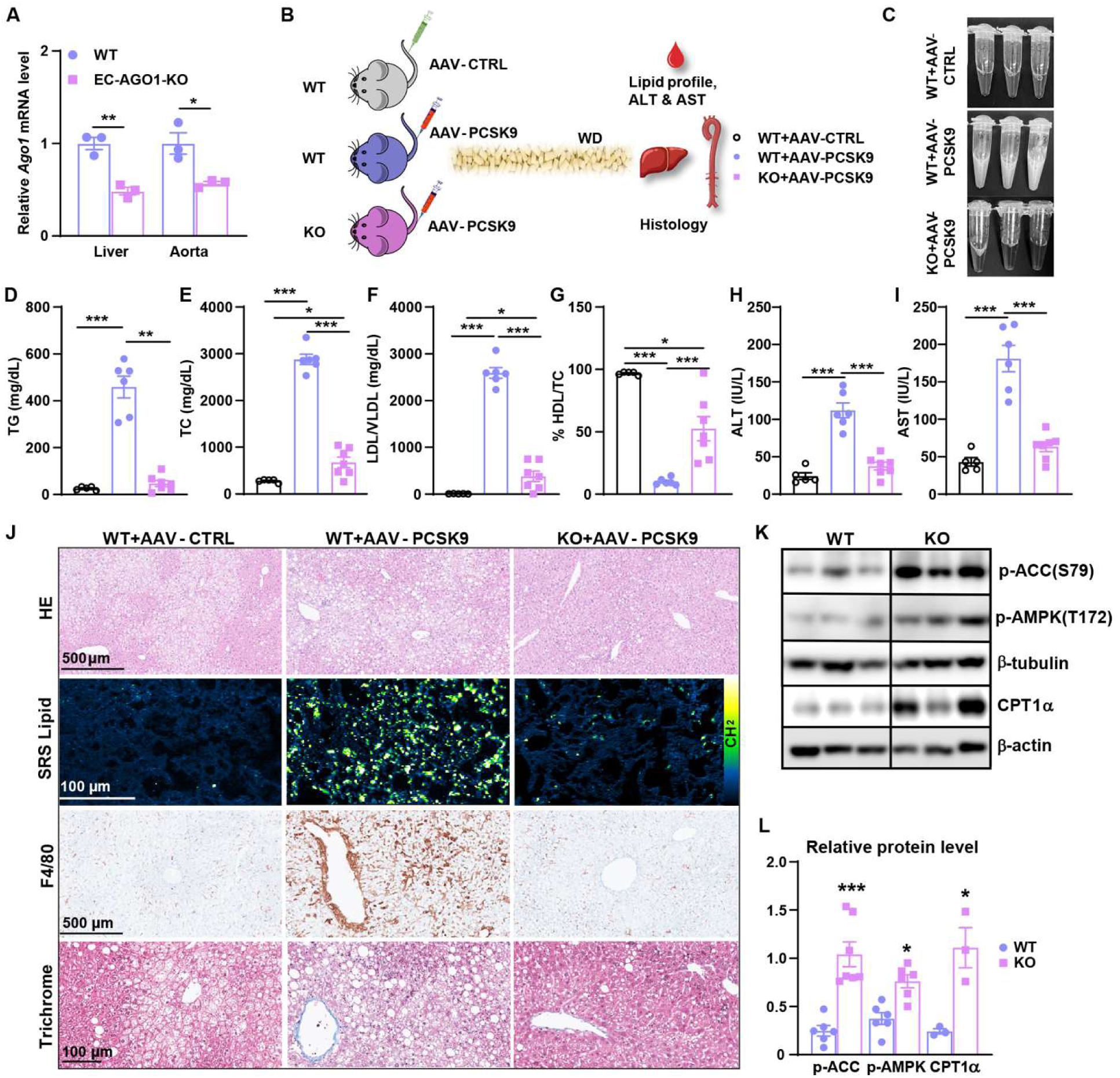
EC-AGO1-KO mice show improved lipid profile and liver function in an AAV9-PCSK9-induced atherosclerosis model. **A**, mRNA levels of AGO1 in ECs isolated from the liver and aorta quantified by qPCR and normalized to 36B4 (n=3 per group). **B-J**, Male EC-AGO1-KO mice and WT littermates (12-week-old) were injected with AAV9-CTRL/PCSK9 and fed a Western Diet for 16 weeks (n=5-7 per group). **B**, Experimental scheme. **C**, Representative plasma samples from three groups of mice described in (**B**). **D-G**, Levels of triglyceride (TG), total cholesterol (TC), LDL/VLDL, and HDL in the plasma after 4 hours fasting. **H**, **I**, Plasma ALT and AST levels. **J**, Representative images of HE, lipid channel (–CH_2_, 2845 cm^-1^) from SRS imaging, F4/80, and Masson’s trichrome staining of the liver. **K**, **L**, Representative images (**K**) and quantification (**H**) of Western blotting for p-ACC1, p-AMPK and CPT1α in livers. Data are presented as mean±SEM in (**A** and **D-I, L**). *p<0.05, **p<0.01, ***p<0.001 between the indicated groups based on Student’s t-test (**A, L**) and one-way ANOVA (**D-I**).

Next, we subjected the WT and KO mice to an atherogenic model induced by AAV9-PCSK9 (**Figure 1B**), which elevates plasma LDL-cholesterol levels by promoting the degradation of hepatic LDL receptors (LDLR).^32^ When combined with a Western-type atherogenic diet (WD; 21% fat, 0.2% cholesterol, ∼34% sucrose), AAV9-PCSK9 induces robust hyperlipidemia and atherosclerosis.^33^ The high sucrose content may accelerate metabolic dysfunction and promote more diffuse aortic lipid deposition, providing a relevant model for studying vascular–metabolic crosstalk.^34,35^ As expected, AAV9-PCSK9 administration increased PCSK9 expression dose-dependently, reaching ∼20-fold induction in mice given 10^11^ vector genomes (vg) (**Figure S3**). WT and KO showed a comparable uptake of AAV9 tracked by tdTomato (**Figure S4**) and PCSK9 protein levels after receiving the same dose of AAV9-PCSK9 (**Figure S5**).

Compared to WT mice given the control AAV (AAV-CTRL) and WD, WT mice receiving AAV9-PCSK9 and WD developed pronounced hyperlipidemia with milky-white plasma, elevated plasma TG, TC, and LDL/VLDL and diminished HDL ratio. In contrast, the EC-AGO1-KO mice subjected to the same regimen showed markedly less hyperlipidemia, hypercholesterolemia, and chylomicronemia (**Figure 1C-G**) without significant differences in body weight (**Figure S6**). In WT mice, the atherogenic regimen increased the plasma alanine aminotransferase (ALT) and aspartate aminotransferase (AST) levels, which were drastically lower in the KO mice, almost comparable to those of WT receiving control AAV (**Figure 1H, I**).

Histological analysis of livers from WT mice subjected to AAV-PCSK9+WD revealed severe steatosis, which was markedly attenuated in EC-AGO1-KO mice. Specifically, KO livers exhibited reduced lipid accumulation, as shown (by HE staining and stimulated Raman scattering [SRS] lipid imaging, along with decreased macrophage infiltration (F4/80 staining) and fibrosis (Masson’s trichrome staining) (**Figure 1J**). KO livers also showed increased levels of phospho-AMPK (p-AMPK), phospho-acetyl-CoA carboxylase (p-ACC), and carnitine palmitoyltransferase 1α (CPT1α), the rate-limiting enzyme for fatty-acid oxidation (FAO), all indicative of enhanced lipid catabolism (**Figure 1K, L**).

### EC-AGO1-KO mice show attenuated atherosclerotic lesions

As expected, *en face* Oil Red O (ORO) staining revealed no obvious plaque in the aortas of the WT mice receiving AAV-CTRL despite the WD diet. This was in stark contrast to the extensive plaque found in the aortic arch, thoracic aorta, abdominal aorta, and aortic roots from WT mice receiving AAV9-PCSK9 and WD. However, the same regimen induced significantly less plaque burden in all vascular regions examined in the EC-AGO1-KO mice (**Figure 2A-D, Figure S7**).

**Figure 2.**
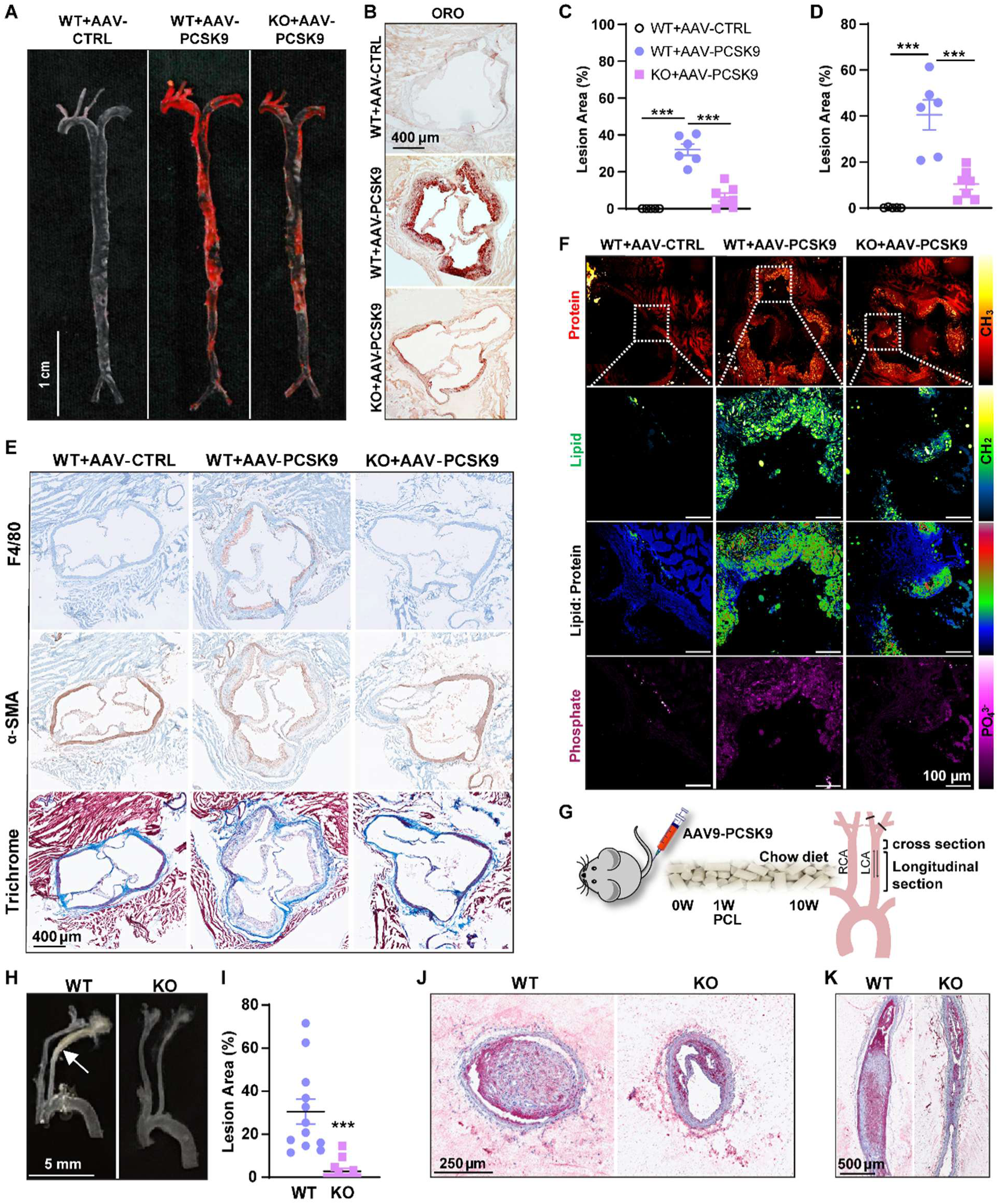
EC-AGO1-KO mice show attenuated aortic atherosclerotic lesion. The same mice described in Figure 1 were analyzed. **A**, **B**, *En face* Oil red O (ORO) staining of the aortic tree and root. **C**, **D**, Quantification of atherosclerotic lesion areas based on ORO staining in the entire aorta (in **C**) and the aortic root (in **D**). **E**, IHC for F4/80, α-SMA, and Masson’s trichrome staining in the aortic root. **F**, SRS imaging of the aortic root. The first row represents a mosaic of the protein channel (CH_3_, 2940 cm^-1^), followed by a zoomed-in image of the lipid channel (CH_2_, 2845 cm^-1^), a corresponding lipid-to-protein ratiometric image, and an image targeted at the phosphate channel (PO_4_^3-^, 960 cm^-1^). Scale bar = 100 µm. Representative images from 5-7 mice/group. **G-K**, Male C57BL/6 WT mice (24-week-old) received AAV9-PCSK9 and were fed a chow diet. One week later, partial carotid artery ligation (PCL) was performed, the ligated left carotid artery (LCA) and the non-ligated right carotid artery (RCA) were harvest at 10 weeks. **H**, **I**, Gross LCA image (**H**) and quantification of lesion area (**I**). **J**, **K**, Representative ORO staining of cross (**J**) and longitudinal sections (**K**) of LCA (as indicated in **G**). Scale bar = 5 mm in **H**, 250 µm in **J** and 500 µm in **K**. Data are presented as mean±SEM (**C**, **D, I**). ***p<0.001 between indicated groups based on one-way ANOVA (**C**, **D**) and Student’s t-test (**I**).

Histological analyses showed that the AAV9-PCSK9-mediated increase in macrophage infiltration (stained by F4/80) and decrease in contractile smooth muscle cells (SMC) content (stained by α-smooth muscle actin [SMA]) and collagen (stained by Masson’s trichrome) were strongly attenuated in the KO mice (**Figure 2E**). To further characterize the biochemical composition of the atherosclerotic lesions, we utilized label-free SRS microscopy which allows non-invasive and high-resolution mapping of metabolites by detecting their vibrational signatures in tissues.^36–38^ In addition to detecting increased lipids in the atherosclerotic plaques from AAV9-PCSK9 treated WT mice, SRS microscopy revealed increased phosphate (PO_4_^3−^, ν_1_ stretching) signals corresponding to the microcalcifications often noted in unstable atherosclerotic plaques.^39^ These metabolite changes were much less in KO mice subjected to the same atherogenic regimen (**Figure 2F**).

The reduced hyperlipidemia observed in the EC-AGO1-KO mice (**Figure 1**) may have contributed to the attenuated atherosclerosis in these animals. To determine whether EC-AGO1 deficiency exerts anti-atherogenic effects independent of lipid lowering, WT and EC-AGO1-KO mice were subjected to AAV-PCSK9 and partial carotid ligation (PCL) under chow diet conditions (**Figure 2G**). Under these conditions, plasma lipid profiles were comparable between groups (**Figure S8**). PCL accelerates atherosclerotic lesion formation through induction of disturbed flow^40^. While all WT mice (n=12) developed robust lesions in the ligated left carotid artery (LCA), only 5 of 12 EC-AGO1-KO mice developed lesions, which were markedly smaller than those observed in WT mice (**Figure 2H-K**). Collectively, these findings indicate that endothelial AGO1 suppression protects against atherosclerosis through both vascular and likely hepatic mechanisms.

### Inhibition of AGO1 in ECs exerts an anti-inflammatory effect

Inflammation links EC dysfunction to atherosclerosis.^3,41,42^ Indeed, in the PCL model, the disturb flow-induced inflammatory markers genes, *Ccl2* and *Thbs1* in the WT LCA were strongly attenuated by EC-AGO1-KO (**Figure 3A**). Consistently, these genes, together with *Icam1* and *Vcam1*, canonical markers of EC inflammation, were decreased in the liver and aorta of EC-AGO1-KO mice subjected to AAV-PCSK9+WD compared with WT littermates. (**Figure 3B, C**).

**Figure 3.**
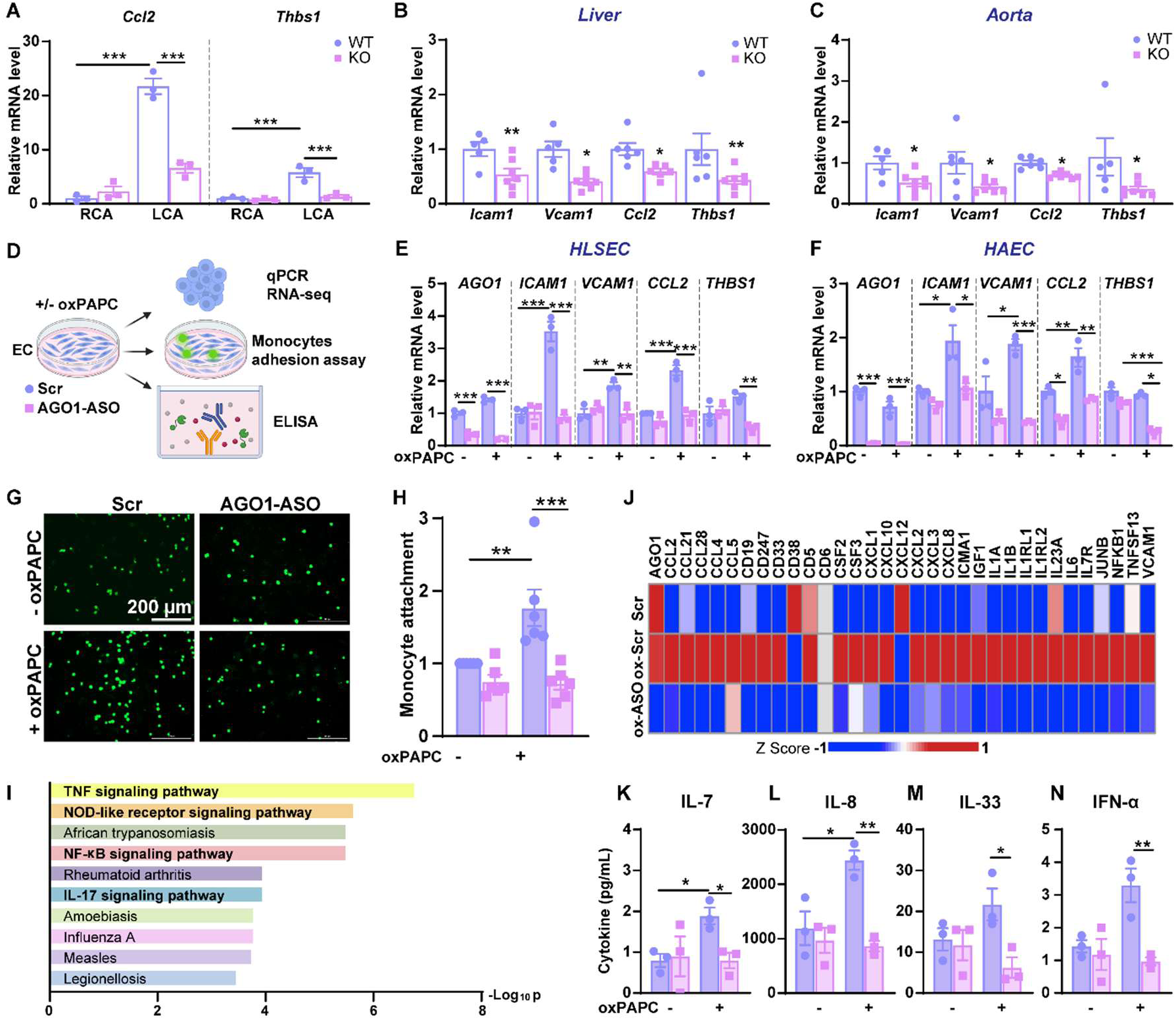
Inhibition of AGO1 in ECs exerts anti-inflammatory effect. **A**, PCL was performed on 12-week-old male EC-AGO1-KO mice and WT littermates fed a chow diet. The LCA and RCA were harvested 1-week post-ligation. mRNA levels *Ccl2* and *Thbs1* were quantified by qPCR and normalized to 36B4. **B**, **C**, qPCR of mRNA levels of inflammatory marker genes in the liver and aorta of mice described in Figure 1. **D**-**N**, ECs were transfected with scramble (Scr) or AGO1-ASO (20 nM), treated with oxPAPC (40 µg/mL) or a vehicle control for 4 hours, and subjected to assays illustrated in (**D**). **E**, **F**, mRNA levels of genes as indicated in human liver sinusoidal ECs (HLSECs) (in **E**) and human aortic ECs (HAECs) (in **F**) were quantified by qPCR, with β-actin as an internal control. **G**, **H**, Attachment of fluorescently labeled THP-1 monocytes to HAECs were imaged and quantified. Shown are relative fold change, with Scr-vehicle control set as 1. Scale bar = 200 µm. **I**, **J**, Bulk RNA-seq of HAECs treated as in **F**. **I**, Enriched KEGG pathways based on KEGG as the effect of AGO1-KD in oxPAPC-treated HAECs. **J**, Heatmap showing induction of genes involved in the cytokine and immune response pathways in HAECs by oxPAPC and downregulation by AGO1-KD. **K-N**, ASO transfected HUVECs were treated with oxPAPC or vehicle control for 4 hours. The media was collected for Luminex ELISA (**K-N**). Data represents mean±SEM from n=3/group (**A**) and n=6-7/group (**B**, **C**) of mice, three (**E**, **F**, **K-N**) and six (**G**, **H**) independent experiments. *p<0.05, **p<0.01, and ***p<0.001 based on two-way ANOVA (**A**, **E**, **F**, and **H**) and Student’s t-test (**B**, **C**).

To determine the effect of AGO1 inhibition in ECs, we designed locked nucleic acid-GapmeR-based antisense oligos (ASOs) that target both human and mouse AGO1 (**Table S1**). Among three ASOs, ASO-112 (termed AGO1-ASO) that showed the strongest and most consistent knockdown (KD) (**Figure S9**) was selected for subsequent experiments. Oxidized 1-palmitoyl-2-arachidonoyl-sn-glycero-3-phosphocholine (oxPAPC), a major component of minimally modified LDL, is known to promote EC dysfunction^43^. In HLSECs and HAECs, AGO1-KD attenuated oxPAPC-induced expression of pro-inflammatory genes, including *ICAM1, VCAM1, CCL2* (encoding monocyte chemoattractant protein 1), and *THBS1* (**Figure 3D-F**). In line with the effect of AGO1-KD on gene expression, oxPAPC-induced monocyte adhesion to HAECs was abolished by AGO1-KD (**Figure 3G, H**). This anti-inflammatory effect of AGO1-KD was also observed in ECs exposed to high glucose and TNF-α (HT), a combination that mimics the metabolic stress induced by the high-sucrose WD and that drives EC dysfunction associated with metabolic syndrome, a major risk factor for atherosclerosis ^44,45^ (**Figure S10**).

To more comprehensively define the effect of AGO1 inhibition on endothelial gene expression, we performed bulk RNA-seq of HAECs following AGO1-KD. Compared to the scramble control (Scr), AGO1-ASO resulted in a total of 195 differentially expressed genes (DEGs) (including 93 up- and 102 down-regulated) in ECs-treated with oxPAPC. The identified DEGs are enriched for immune and inflammatory response pathways, e.g., TNF-α signaling, NOD-like receptor signaling, and NF-κB signaling (**Figure 3I**). A panel of oxPAPC-induced inflammatory genes, including adhesion molecules (*ICAM1* and *VCAM1*), chemokine ligands (i.e., C-C motif chemokine ligands [CCLs] and C-X-C motif chemokine ligands [CXCLs]), and interleukins (ILs), were suppressed by AGO1-KD (**Figure 3J**). In parallel, we analyzed the cytokine production from ECs using the Luminex assay (**Figure 3D**). At baseline, AGO1-KD did not significantly affect the levels of most of the cytokines assayed. However, in ECs treated with oxPAPC, AGO1-KD significantly reduced the levels of several pro-inflammatory cytokines including IL-7, IL-8, IL-33, and IFN-α (**Figure 3D, K-N**). These protein changes were consistent with the transcript changes (**Figures 3J and S11**).

### AGO1 localizes to the nucleus and interacts with NF-κB to drive pro-inflammatory gene expression in ECs

The strong anti-inflammatory effect and the decreased mRNA levels of pro-inflammatory genes due to AGO1 KO/KD in ECs suggest a positive regulation of inflammation by AGO1, likely through transcriptional regulation. In cancer cells, AGO1 was shown to bind to promoters and interact with RNA polymerase II (RNAP II) and TFs such as estrogen receptor (ER) to facilitate gene activation.^46,47^ To determine whether a similar if such a mechanism exists in non-cancerous cells like ECs, we first performed immunofluorescent (IF) staining of AGO1. We detected AGO1 in the nucleus and cytoplasm in ECs at baseline and under inflammatory stress without significant difference in either protein levels or subcellular localizations (**Figure 4A, B, Figure S12A-D**). This was consistent with Western blotting of AGO1 in subcellular fractionations from ECs (**Figure 4C, Figure S12E**). To investigate the function of nuclear AGO1 in ECs, we performed Cut&Tag-sequencing (Cut&Tag-seq) using an AGO1 antibody, with IgG as an isotype control. Cut&Tag revealed strong binding between AGO1 and chromatin, which was increased by oxPAPC but decreased by AGO1-KD in ECs (**Figure 4D**). In oxPAPC-treated ECs, gene promoters accounted for 24% of AGO1-bound genomic regions, which was reduced to 19% by AGO1-KD (**Figure 4E**).

**Figure 4.**
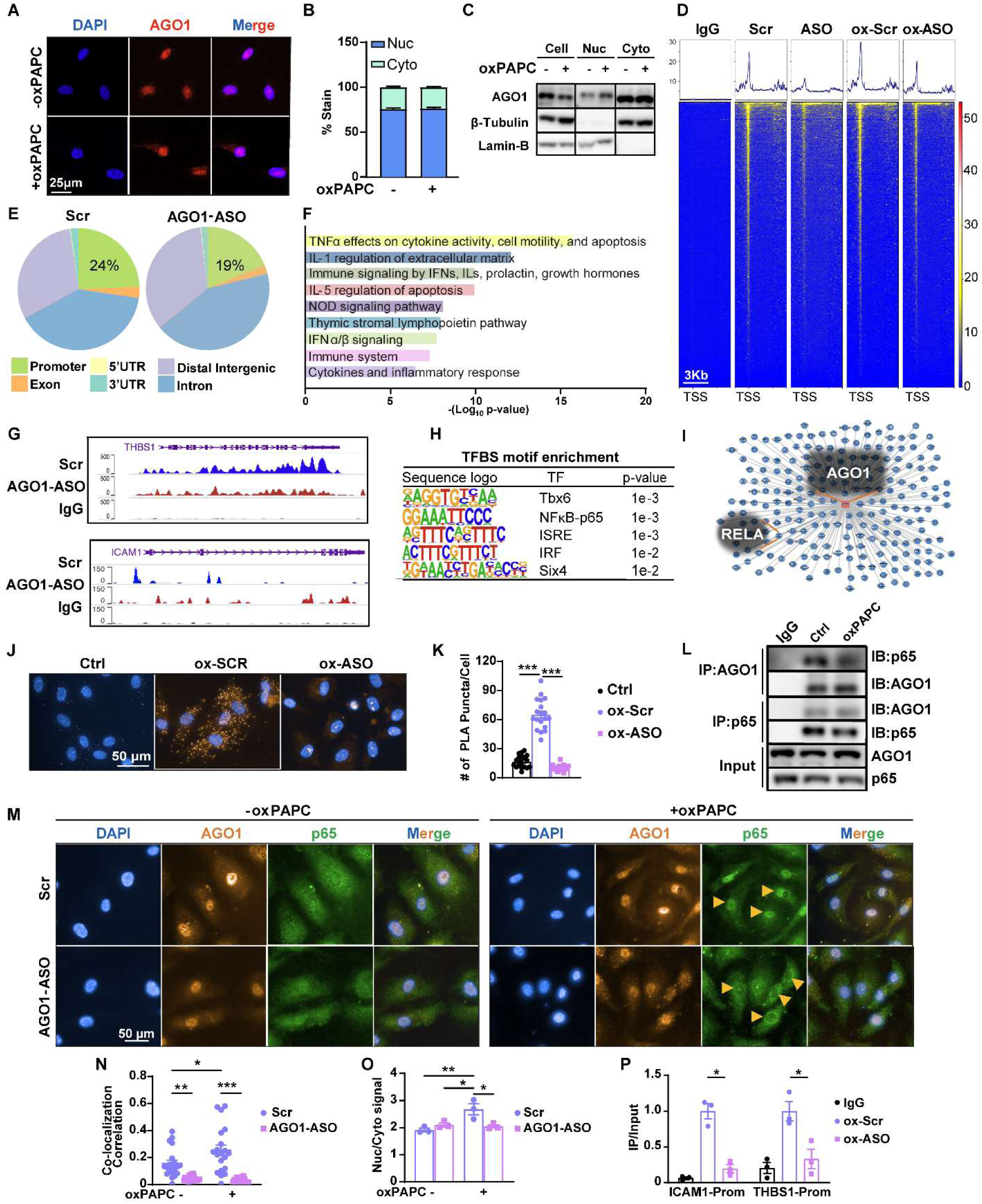
AGO1 interacts with NF-kB to regulate the pro-inflammatory gene expression. A-C, HAECs were treated with vehicle control or oxPAPC for 4 hr. Confocal imaging of AGO1 immunofluorescent (IF) staining was performed with DAPI counterstain (**A**) and the subcellular localization was quantified (**B**). Scale bar = 25 µm. **C**, Representative Western blot of AGO1 in subcellular fractions, β-tubulin and Lamin-B1 were used as cytoplasmic (cyto) and nuclear (nuc) markers. **D**, **E**, HUVECs transfected by ASO and treated with or without oxPAPC underwent Cut&Tag-seq with AGO1 antibody, with IgG as an isotype control. **D**, AGO1 occupancy in the chromatin, with enriched peaks near transcription start sites (TSS). **E**, Distribution of AGO1-binding peaks across the promoter and other genomic regions in oxPAPC-treated ECs without or with AGO1-KD. **F**, Enriched pathways in the 86 genes that are regulated by AGO1 and show promoter binding by AGO1. **G**, Gene tracks showing the binding of AGO1 in the genomic loci encoding indicated marker genes. **H**, Top 5 TFs (including NF-κB RELA) whose TFBS were enriched in AGO1-bound DNA sequences identified by Cut&Tag. **I**, AGO1 protein-protein interaction network constructed based on yeast-two hybridization. **J**, **K**, **M-P**, HAECs were transfected with ASO and treated with oxPAPC or vehicle control for 4 hours, AGO1-p65 interactions was detected by proximity ligation assay (PLA). Representative images (**J**) and quantifications (**K**) are shown. Scale bar = 50 µm. **L**, HUVECs were treated with or without oxPAPC. Co-IP of AGO1 and NF-κB p65, with IgG as an isotype control. **M-O**, Representative images of co-IF of AGO1 and p65 in HAECs as in J, K (**M**). Scale bar = 50 µm. **N**, Quantification of nuclear co-localization of AGO1 and p65. **O**, Quantification of the p65 localization. **P**, HAECs were treated with oxPAPC without or with AGO1-KD. ChIP was performed with p65 antibody and the association of p65 with *THBS1* and *ICAM* promoters was detected using qPCR. Data are presented as mean±SEM, *p<0.05, **p<0.01 and ***p<0.001 based on one-way ANOVA (**K**). and two-way ANOVA (**N-P**).

Combining the RNA-seq (**Figure 3I, J**) and Cut&Tag-seq datasets, we found that out of 102 genes downregulated by AGO1-KD in oxPAPC-treated ECs, 86 showed association with AGO1 occupancy in their promoter regions. Of these, 60 are involved in the endothelial immune or inflammatory responses of ECs (**Figure 4F**), including *THBS1* and *ICAM1* (**Figure 4G**). As validation, qPCR revealed that AGO1 indeed binds to the peak regions in the promoters of *THBS1* and *ICAM1* as identified by Cut&Tag-seq and AGO1-KD decreased these interactions in oxPAPC-treated ECs (**Figure S13**).

In breast cancer cells, AGO1 was found to act as a transcriptional co-activator of ER alpha.^47^ To identify potential TFs involved in AGO1-regulated inflammatory gene program in ECs, we analyzed TF binding sites (TFBS) in the promoter regions of 86 genes that are both AGO1-bound and AGO1-positively regulated. Using TRANSFAC, NF-κB (subunit p65, aka RELA) emerged as one of the top candidates (**Figure 4H**). Out of the 86 genes identified, 17 have been previously reported to be regulated by NF-κB with putative p65 TFBS^48,49^, including *THBS1* and *ICAM1* (**Table S2**). Additionally, NF-κB signaling was among the top enriched pathways in DEGs caused by AGO1-KD (**Figure 3I**). Search using databases for protein-protein interactions (PINA https://omics.bjcancer.org/pina and INTACT https://www.ebi.ac.uk/intact) revealed that AGO1-RELA interaction has been experimentally identified by using a yeast-two hybrid system^50^ (**Figure 4I**).

In ECs, proximity ligation assay (PLA) detected AGO1-p65 interactions in oxPAPC-treated ECs, which signal was reduced by AGO1-KD (**Figure 4J, K**). The AGO1-p65 interaction was also confirmed by co-immunoprecipitation (co-IP) (**Figure 4L**) and co-immunofluorescence (co-IF) (**Figure 4M, N**, **Figure S14**). Importantly, Co-IF also revealed that AGO1-KD substantially decreased p65 nuclear localization (**Figure 4M, O**), although IκBα, whose degradation is key to p65 activation,^51,52^ was unaltered (**Figure S15**). Furthermore, p65 binding to the *ICAM1* and *THBS1* promoters was reduced by AGO1-KD (**Figure 4P**). Together, these results suggest that AGO1 binds to NF-κB and promotes its nuclear translocation and transactivation of pro-inflammatory genes in ECs.

### Lipid nanoparticle (LNP) delivery of AGO1 ASO confers anti-hyperlipidemic and anti-atherosclerotic effects

Given that EC-AGO1-KO mice showed anti-hyperlipidemic and anti-atherosclerotic tendencies, we evaluated the therapeutic potential of AGO1-ASO to treat atherosclerosis. We confirmed the efficacy of AGO1-ASO in WT mice. Intravenous injection (i.v.) of AGO1-ASO (10 mg/kg body weight) decreased *Ago1* mRNA levels in the liver of WT mice by more than 50% (**Figure S16A**). When encapsulated in LNP, AGO1-ASO (1 mg/kg body weight, i.e., 1/10^th^ of the dose without LNP) reduced *AGO1* mRNA by 75% (**Figure S16B**) and provided a dose for further study. As expected,^53^ biodistribution analysis confirmed that LNP were predominantly accumulated in liver, with lower signals detected in the aorta, heart, spleen, kidney, lung and skeletal muscle (**Figure S17**).

WT mice were given AAV9-PCSK9 and WD. Two weeks later, mice received weekly i.v. injections of LNP-AGO1-ASO or LNP-Scr for 4 weeks (**Figure 5A**). LNP-AGO1-ASO-treated mice had almost clear plasma (**Figure 5B**), along with significantly reduced TG, TC, and LDL/VLDL levels, and an increased HDL/TC ratio (**Figure 5C-F**). At the gene expression level, LNP-AGO1-ASO reduced *Ago1* expression in both the liver and aorta. In the liver, this was accompanied by decreased expression of pro-inflammatory (*Icam1, Vcam1, Ccl2, Thbs1*) and pro-fibrotic (*Acta2, Collagen I, Collagen III*) genes (**Figure 5G**). A similar anti-inflammatory effect was observed in the aorta (**Figure 5H**).

**Figure 5.**
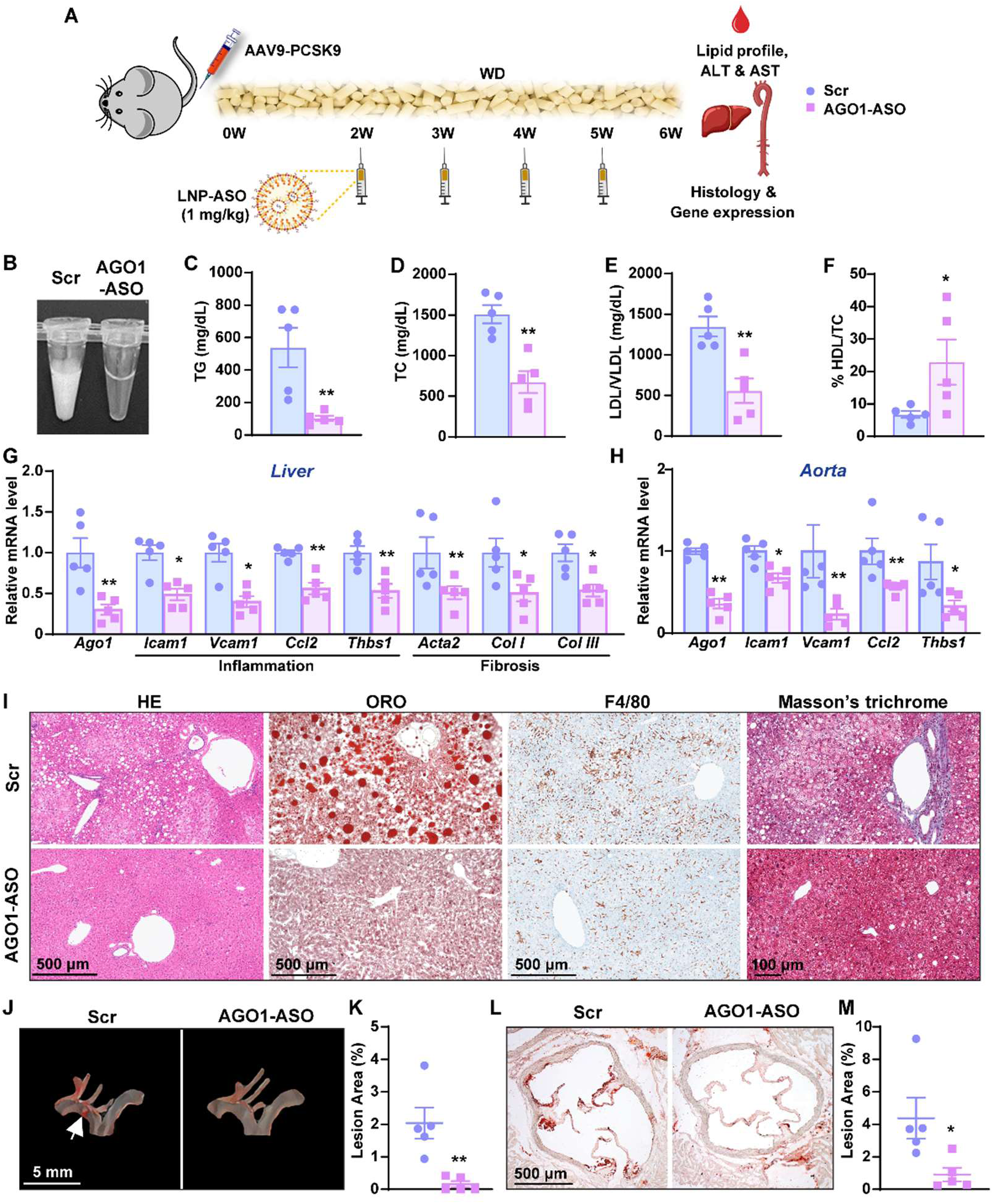
Lipid nanoparticle (LNP) delivery of AGO1 ASO is anti-hyperlipidemic and anti-atherosclerotic. **A**, Experimental design: male C57BL6 wildtype mice (20-week-old, n=5/group) received AAV9-PCSK9 and were fed a WD. Two weeks later, LNP-ASO was injected through tail vein at 1 mg/kg body weight weekly for 4 weeks. **B**, Plasma samples were collected from LNP-ASO treated mice after 4 hours of fasting. **C-F**, Plasma lipid profiles. **G**, **H**, qPCR quantification of mRNA levels of various genes in the liver (in **G**) and aorta (in **H**), including aortic EC Ago1 expression. **I**, Representative images of HE, ORO, F4/80 and Masson’s trichrome staining of the liver. **J-M**, Representative images of ORO staining of the aortic arch and root and the quantification. Data are presented as mean±SEM, *p<0.05, **p<0.01 based on Student’s t-test (**C-H**, **K** and **M**).

LNP-AGO1-ASO-treated mice also had significantly less hepatic lipid accumulation, macrophage infiltration, and fibrosis (**Figure 5I**). Importantly, LNP-AGO1-ASO treatment did not affect liver size, or plasma ALT and AST levels (**Figure S18**), suggesting it was not toxic to the liver. Finally, the atherosclerotic lesions in the aortic arch and root were substantially attenuated in LNP-AGO1-ASO-treated mice (**Figure 5J-M**). These findings demonstrate that systemic AGO1 inhibition is anti-hyperlipidemic and anti-atherosclerotic.

### Targeted delivery of AGO1 ASO to inflamed endothelial cells via monocyte membrane–coated nanoparticle (MoNP) reduces hyperlipidemia and atherosclerosis

To evaluate the therapeutic effect of EC-AGO1 targeting in an established atherosclerosis model, we leveraged a biomimetic MoNP platform. We showed that monocyte membrane cloaking enhances the preferential targeting of NP to regions of vascular inflammation, enabling effective delivery of a small-molecule inhibitor drug to inflamed ECs and attenuation of atherosclerotic lesion.^11^ We adapted MoNP to deliver AGO1-ASO (**Figure S19A-C**) to mice with established atherosclerotic plaques. The MoNPs were characterized and found to reduce *Ago1* expression in HAECs (**Figure S19D**). In addition, MoNP exhibited endothelial homing both in liver and aorta of the atherosclerotic mice (**Figure S20**).

We first induced advanced atherosclerosis in the WT mice by using AAV-PCSK9 and WD feeding for 5 months, followed by weekly i.v. injections of MoNP-AGO1-ASO or MoNP-Scr at 1 mg/kg for 4 weeks (**Figure 6A**). Compared with MoNP-Scr-treated controls, MoNP-AGO1-ASO-treated mice exhibited visibly clearer plasma (**Figure 6B**) and improved lipid profiles (**Figure 6C-F**). As expected, AGO1 expression levels were decreased in liver ECs (**Figure 6G**) and marker genes of inflammation and fibrosis were also decreased in the liver and aortic ECs (**Figure 6H, I**). MoNP-AGO1-ASO treatment also improved liver histology (**Figure 6J**), without altering ALT and AST levels (**Figure S21**). Furthermore, MoNP-AGO1-ASO significantly reduced atherosclerotic burden in the aortic arch (**Figure 6K, L**). These effects largely recapitulated those observed in mice receiving LNP-delivered AGO1-ASO and EC-AGO1-KO mice.

**Figure 6.**
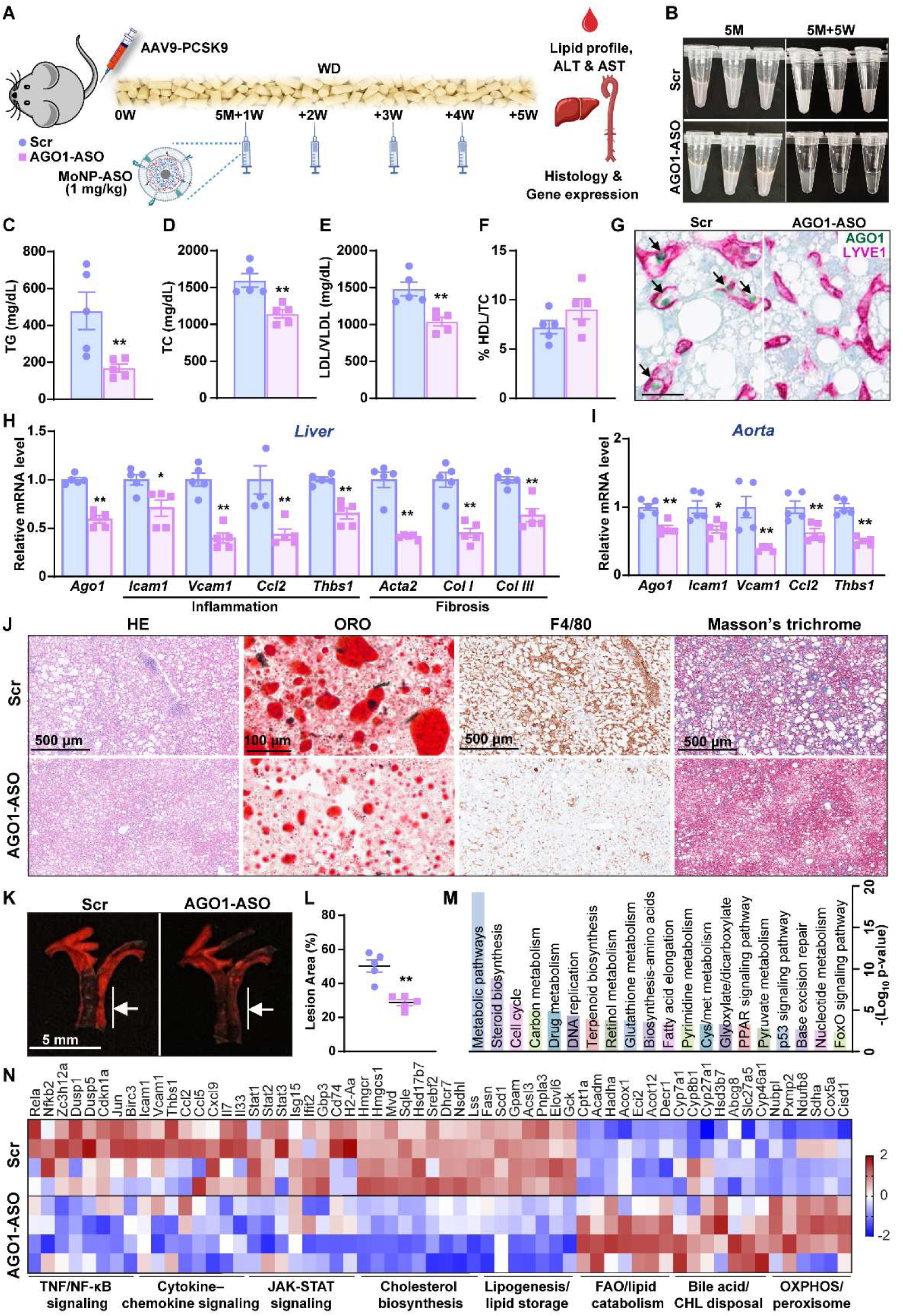
Monocyte membrane-coated nanoparticles (MoNP) delivery of AGO1-ASO is anti-hyperlipidemic and anti-atherosclerotic. **A**, Experimental design: male C57BL6 WT mice (20-week-old, n=5/group) received AAV-PCSK9 and were fed a WD. Five months later, MoNP-ASO was given i.v. at 1 mg/kg body weight weekly for 4 weeks. **B**, Appearance of plasma samples collected before and after MoNP-ASO treatment. **C-F**, Plasma lipid profiles. **G**, Representative images of AGO1 (green) and LYVE1 (pink) dual IHC staining of liver. Scale bar = 25 μm. **H**, **I**, qPCR quantification of mRNA levels of various genes in the liver (in **H**) and aortic EC (in **I**). **J**, Representative images of HE, ORO, F4/80 and Masson’s trichrome staining of the liver. **K-L**, Representative images and quantification of ORO staining of the aortic arch. **M**, **N**, Bulk RNA-seq of livers. **M**, Enriched pathways based on KEGG as the effect of MoNP-AGO1-ASO in livers. **N**, Heatmap showing changes of genes involved in selected pathways comparing between MoNP-Scr vs MoNP-AGO1-ASO treated livers. Data are presented as mean±SEM, *p<0.05, **p<0.01 based on Student’s t-test (**C-F**, **H**, **I**, and **L**).

To gain more understanding of the effect of MoNP-AGO1-ASO, we performed RNA-seq of the liver. Compared to the MoNP-scramble control, MoNP-AGO1-ASO led to 368 DEGs, which were enriched in metabolic pathways, steroid biosynthesis, cell cycle pathways, etc. Lipogenesis and cholesterol biosynthesis pathways were downregulated, whereas FAO, cholesterol disposal, and mitochondrial/peroxisomal metabolic pathways were upregulated. These findings were consistent with increased expression of CPT1α in EC-AGO1-KO mice (**Figure 6M, N**). These findings, together with those from LNP-AGO1-ASO, demonstrate that targeting AGO1 using either systemic or EC-targeting strategies attenuates hyperlipidemia and atherosclerosis.

## DISCUSSION

This study points out a critical role of AGO1 in driving EC dysfunction, vascular inflammation, and liver dysfunction in the context of atherosclerosis (**Figure 7**). EC-AGO1-KO mice exposed to AAV-PCSK9 and an atherogenic diet exhibited an improved lipid profile and liver function, decreased hepatic steatosis, inflammation, and fibrosis, and substantially attenuated atherosclerotic lesions. Likewise, LNP-AGO1-ASO and MoNP-AGO1-ASO treated WT mice resisted the metabolic and vascular derangements induced by PCSK9 and an atherogenic diet. Together with our previous findings showing an anti-obesity and anti-insulin resistance phenotype in EC-AGO1-KO mice subjected to a HFHS diet,^27^ the pleiotropic effects of EC-AGO1 inhibition underscore the importance of EC dysfunction in both metabolic disorders and vascular diseases.

**Figure 7.**
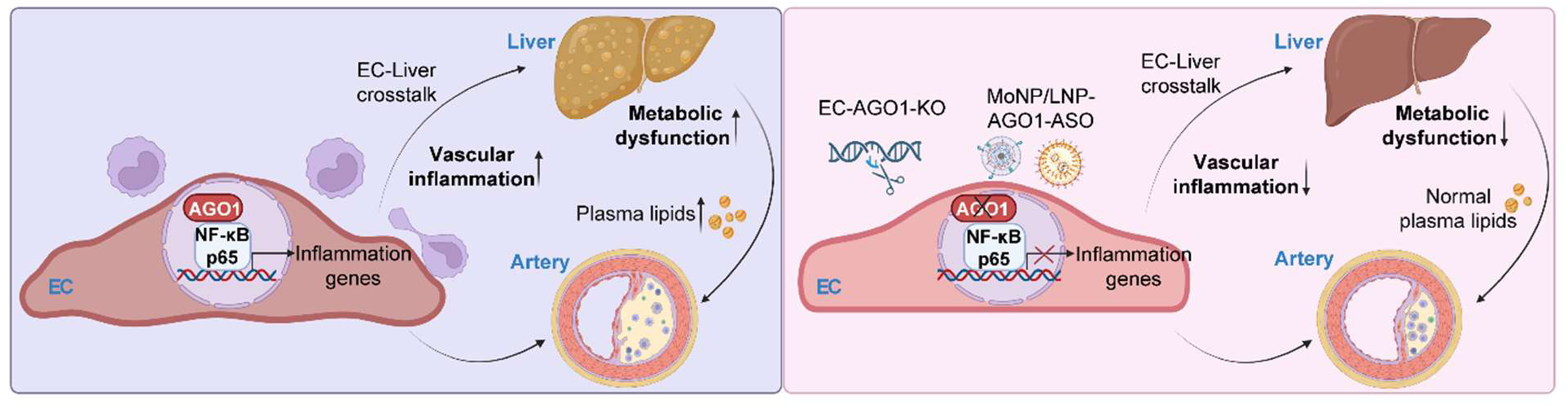
Schematic summary. Schematic summary of the proposed mechanism of AGO1-regulated inflammatory response in ECs in the context of hyperlipidemia and atherosclerosis (left) and the effect of EC-AGO1 inhibition on vascular inflammation, liver dysfunction, and atherosclerosis (right). Image credit: Figure created with BioRender.

The benefits of EC-AGO1-KO and inhibition, as shown *in vitro* and *in vivo*, indicate a consistent anti-inflammatory effect in the liver and in the aortas/arteries. Remarkably, these effects were mostly observed in ECs and animals exposed to inflammatory and metabolic stress, but not at baseline, suggesting a context-dependent aspect to AGO1 KO or inhibition in ECs. The strong suppressive effect on hypercholesterolemia in mice with EC-AGO1 inhibition, the reduced atherosclerotic burden could be due to improved lipid profile, rather than reduced inflammation in the arteries. To specifically address the effect of EC-AGO1-KO on the arterial inflammation, we use the PCL models and observed decreased inflammatory gene expression and atherosclerotic lesion formation. Thus, EC-AGO1 inhibition likely exerts anti-inflammatory effects in both liver and blood vessels, together contributing to an anti-atherosclerotic phenotype.

Regarding the effect of endothelial AGO1 suppression on the liver, we identified a strong anti-inflammatory program. AGO1-KD in oxPAPC-treated ECs suppressed expression of adhesion molecules (*ICAM1* and *VCAM1*), inflammatory factors/cytokines (*THBS1*, *IL1A*, and *IL1B*) and chemokines (*CCL2*, *CXCL1*, and *CXCL2*), and production of IL-7, IL-8, IL-33, and IFN-α, all of which can promote hepatocyte dysfunction.^54–57^ These molecular changes are consistent with reduced monocyte adhesion to ECs with AGO1-KD and reduced hepatic macrophage infiltration in EC-AGO1-KO mice, pointing to EC-AGO1 orchestrating hepatic inflammation.

ECs also regulate hepatocyte metabolism and function through paracrine signaling.^15,17,18,58^ In a EC-HepG2 co-culture system, AGO1-KD in ECs caused a decrease in inflammatory gene expression (*ICAM1*, *CCL2* and *THBS1*), but an overall increase of genes involved in lipid metabolism, including fatty acid uptake and oxidation and cholesterol degradation, as well as bile acid synthesis in HepG2 cells (**Figure S22A-C**). These data suggest that EC-derived paracrine signals may suppress inflammation while enhancing the metabolic function of hepatocytes.

Wnt/β-catenin signaling regulates hepatocyte cholesterol uptake, bile acid conjugation^15^ and liver zonation^59^. Germane to this, EC-AGO1-KO mice expressed higher levels of *Wnt2* and its downstream *Axin2*, *Tbx3*, *Glul* in the liver (**Figure S22F**). Accordingly, periportal Zone 1, marked by CYP2F2 and associated with oxidative and catabolic metabolism was expanded whereas pericentral Zone 3, marked by glutamine synthase (GS) and associated with glycolysis and lipogenesis was reduced in EC-AGO1-KO **(Figure S22G-J**). Class 3 semaphorins (SEMA3), such as SEMA3A, are produced by LSECs, increased under HFD, and promotes hepatic lipid accumulation possibly through less LSEC fenestration and impaired EC-hepatocyte exchange.^60^ Although AGO1-KD did not markedly alter EC SEMA3A, it significantly reduced SEMA3C expression in oxPAPC-treated and HepG2-cocultured ECs, a finding of potential relevance given the reported role of SEMA3C in promoting liver fibrosis^61^ (**Figure S22K**). Nonetheless, detailed mechanisms linking endothelial AGO1 inhibition to the broad metabolic and functional changes observed in the liver warrant further investigation.

AGO1 has been studied in the context of miR-mediated gene silencing. We previously demonstrated that AGO1 regulates the *THBS1* expression through miR-mediated post-transcriptional silencing, which may contribute to EC dysfunction, obesity, and insulin resistance.^27^ As expected, *THBS1* expression was consistently decreased by AGO1-KD/KO in oxPAPC-treated ECs and in the mouse liver and the aortic tree. Given the reported role of TSP1 in EC dysfunction, inflammation, and fibrosis in the context of MASH and atherosclerosis,^62,63^ inhibition of *THBS1* probably contributes to the beneficial effects of EC-AGO1 inhibition. Aside from this mechanism and the canonical role of AGO1 in RISC, we identified a non-canonical mechanism by which AGO1 interacts with NF-κB (a master regulator of inflammation^52^) and genomic loci encoding key inflammatory genes in the nucleus of ECs. Of note, neither p65/RELA nor its upstream IkBα expression was affected by AGO1-KD. However, AGO1-KD decreased p65 nuclear localization and binding to the promoters of its transcriptional targets. Studies conducted with cancer cells have shown AGO1 was necessary for ER binding to the promoters of its target genes upon estradiol treatment^47^ and AGO1 may play a role in transcriptional activation.^46,64,65^ It is plausible to propose that AGO1 may act as a co-activator for NF-κB-transactivated gene expression. Given the case of *THBS1*, such nuclear action of AGO1 is likely in synergy with its cytoplasmic function in mediating RISC, collectively contributing to EC dysfunction under pro-inflammatory conditions. Future studies are warranted to further define the molecular basis of AGO1-p65 interactions and how this interaction drives inflammatory gene activation.

In the treatment studies, both LNP and MoNP-delivered-AGO1-ASO improved lipid profiles and attenuated atherosclerotic burden without evidence of liver toxicity, largely resembling the phenotypes observed in the EC-AGO1-KO mice. This suggests common underlying mechanisms mediated by EC-AGO1. The two interventions with ASO were evaluated in different stages of atherosclerosis. LNP-AGO1-ASO was tested during lesion formation and demonstrated systemic AGO1 inhibition can suppress atherogenic development. In contrast, MoNP-AGO1-ASO was evaluated in mice with established atherosclerosis, a setting more relevant to the clinical setting.

Notably, MoNP-AGO1-ASO significantly reduced lesion burden despite prolonged WD feeding, underscoring the therapeutic potential of EC-directed AGO1 inhibition even in established atherosclerosis. Of note, AGO1-KD in the hepatocytes did not lead to similar changes as those noted with EC-AGO1-KD (**Figure S22D, E**). Moreover, hepatocyte-specific AGO1-KO mice did not show significant difference in liver function or whole-body metabolism either at baseline or under a high-fat diet.^66^ Collectively, these findings emphasize ECs as major contributors to the anti-hyperlipidemic and anti-atherosclerotic effects of AGO1-ASO. Our data and reports from others^19,27,28,67^ support the view that EC-directed therapeutics may simultaneously improve metabolic and cardiovascular outcomes.^31^

Sex-dependent differences in experimental atherosclerosis models are well recognized and female mice are generally more resistant to diet-induced hyperlipidemia and atherosclerosis.^68,69^ Indeed, the atherogenic regimen used in male mice induced only moderate hyperlipidemia in female mice. Nonetheless, female EC-AGO1-KO mice showed a trend toward decreased LDL/VLDL levels (**Figure S23A-E**) and female mice treated with AGO1-ASO showed lower expression of inflammatory markers (*Icam1* and *Ccl2*) in the liver and aorta, compared with controls (**Figure S23F-H**). Furthermore, when challenged with AAV-PCSK9+WD combined with PCL, WT female mice developed clear atherosclerotic lesions in the ligated carotid artery, whereas EC-AGO1-KO females exhibited marked plaque reduction (**Figure S23I-M**). Together, these findings suggest that the atheroprotective effect of EC-AGO1 deficiency emcompasses both male and female animals.

RNA-based therapeutics are rapid advancing for human diseases.^13,70^ Application of this technology to AGO1 suggests a means of ameliorating EC dysfunction and vascular inflammation in the liver and arterial wall, conferring a protective effect against hypercholesterolemia and its associated atherosclerosis. Inhibition of EC-AGO1 via LNP or MoNP provide strategies to modulate inflamed endothelium and confer vascular and metabolic protection. By enabling EC-targeted RNA delivery, MoNP-based inhibition of EC-AGO1 could tackle metabolic and blood vessel disease while limiting toxicity.

## METHODS

### Study Approval

All animal experiments conducted were approved by the Institutional Animal Care and Use Committees at City of Hope (#17010) and Institutional Biosafety Committee (#16023).

### Mouse Models

EC-AGO1-KO and their WT littermates were generated by crossbreeding VE-Cadherin-Cre (B6.FVB-Tg [Cdh5-cre]7 Mlia/J) and AGO1^flox/flox^ (Ago1tm1.1Tara/J) mice, both with C57BL6 at City of Hope as described.^27^ The WT and KO littermates from the same breeders were housed in the same cages until subjected to metabolic phenotyping and tissue collection. At end points of each experiment, mice were euthanized with CO_2_ inhalation.

### AAV9-PCSK9 model

Mice were injected in the tail vein with AAV-CTRL or recombinant AAV serotype-9 expressing the PCSK9 mutant under the hepatic control region-apolipoprotein enhancer/alpha1-antitrypsin, a liver-specific promoter (AAV9-HCRApoE/hAAT-D377Y-mPCSK9, abbreviated as AAV-PCSK9)^13^ at 1×10^11^ vg and fed a chow diet or a Western-type atherogenic diet (WD; 21% fat, 0.2% cholesterol, ∼34% sucrose by weight) (Harlan-Envigo TD.88137).

### Partial carotid ligation (PCL) model

PCL surgery was performed as published.^40,71^ The left external carotid artery (CA), left internal CA, and left occipital artery were ligated, while the left superior thyroid artery was not.

### ASO design and synthesis, LNP-ASO and MoNP-ASO formulation

#### ASO design and synthesis

The ASOs were designed based on the homologous regions of human and mouse AGO1 mRNA transcripts. ASOs are 16 nt-long gapmers chemically modified for nuclease resistance using fully phosphorothioated backbone and locked nucleic acid (LNA) modification of 1-3 nt at both 5’ and 3’ ends.^72^ Sequences are provided in Table S1. ASO gapmers were synthesized and at the DNA/RNA Synthesis Laboratory (City of Hope) and were purified and analyzed using anion-exchange HPLC, desalted and lyophilized. A scrambled control LNA Gapmer was purchased from QIAGEN with a sequence of 5’-AACACGTCTATACGC-3’ (Product #339516).

#### LNP-ASO formulation

The lipid nanoparticles were synthesized using a microfluidic device (NanoAssemblr Ignite, Precision NanoSystems). The formulation utilized the SM-102 ionizable lipid used in Moderna’s mRNA1273 COVID-19 mRNA vaccine. All lipids, including SM102 (Echelon Bioscience, Cat#N1102) Cholesterol (Avanti Polar Lipids, Cat#700000), DSPC (Avanti Polar Lipids, Cat#850365), and DMG-PEG (Avanti Polar Lipids, Cat#880151), were dissolved in ethanol and mixed at a mole ratio of 50:38.5:10:1.5, respectively. The AGO1-ASO was dissolved in citrate buffer (pH-4, 50 mM). The aqueous and ethanol phases were mixed at a volumetric ratio of 3:1 using a microfluidic device. The formulated LNP-ASO were then dialyzed in 1×PBS using a dialysis bag with a molecular weight cutoff set at of 12-14kDa (Spectra/Por, Cart# 132703T) for 2 hours, then analyzed for encapsulation efficiency, size (97.64 nm), and zeta potential (−0.5 mV).

#### LNP biodistribution in vivo

LNP containing luciferase mRNA (Luc-mRNA-LNP) were injected through i.v. into mice at the dosage of 0.3 mg/kg, control mice were injected PBS (27-week-old, n=5). 24 hours later, plasmas were collected for PBMC isolation. Other tissues were collected for qPCR or IHC.

#### MoNP-AGO1-ASO synthesis and characterization

Ten milligrams of PLGA (Sigma Aldrich) was dissolved in 1 mL acetone to prepare NP cores using a single-emulsion solvent evaporation method. For fluorescence tracking, lipophilic dye DiD was incorporated into the organic phase to generate DiD-loaded NPs. Unlabeled NP cores were coated with branched polyethyleneimine (bPEI, average Mw ∼25,000, Sigma Aldrich) at a 1:1 mass ratio to generate NP-bPEI, followed by loading with AGO1-ASO through electrostatic interactions. The resulting NP-bPEI-AGO1-ASO were confirmed using an agarose gel retardation assay, and the AGO1-ASO loading efficiency was determined by measuring unbound AGO1-ASO in the supernatant using a Qubit assay. Plasma membranes isolated from mouse bone marrow-derived monocytes were used to cloak NP-bPEI-AGO1-ASO following a previously reported method,^11^ yielding MoNP-AGO1-ASO. DiD-loaded MoNPs were prepared using the same membrane-cloaking procedure. Hydrodynamic diameter and zeta-potential of all nanoparticle formulations were measured by dynamic light scattering using a zetasizer (Malvern).

#### MoNP-AGO1-ASO characterization in vivo

C57BL/6J mice were administered with 1×10^11^ vg of AAV8-D377Y-mPCSK9 (Vector Biolabs) or saline and maintained on a high-fat diet (Envigo TD.88137) for 3 weeks. Mice then received 300 μg of DiD-loaded MoNP via intravenous injection. Four hours after MoNP administration, mice were euthanized, and arterial tissues (aorta and carotid arteries) and livers were harvested for IVIS imaging to assess MoNP distribution. Liver and aortic arch tissues were sectioned and immunostained with antibodies against LYVE1 (Cell Signaling Technology, Cat# 67538S, 1:200), and CD31 (BD Biosciences, Cat# 553370, 1:200). Images were captured by BioTek Lionheart LX automated microscope.

Additional material and methods are detailed in the Supplemental Material and Methods.

### Statistics

All *in vitro* data represent at least three independent experiments unless otherwise indicated. All *in vivo* data represent experiments performed with numbers of mice as specified in the figure legends. Statistical analyses for data other than high-throughput sequencing (see Methods in the Data Supplement) were performed using GraphPad Prism. Two-group comparisons were performed using 2-sided Student’s t-test and multiple-group comparisons were performed using ANOVA followed by Tukey’s post-hoc test. *P* values less than 0.05 were considered statistically significant.

## Data availability

All sequencing data are available at GEO numbers for Cut&Tag: GSE293927 and bulk RNA-seq: GSE293932.

## Acknowledgments

The authors wish to thank Dr. Yun Fang at University of Chicago, for the generous gift of AAV9-PCSK9; Drs. Zhao Wang, Holly Yin, and Jiawei Sun and Ms. Muxi Chen at the City of Hope, Dr. Samar Ibrahim at Mayo Clinic, Dr. Oren Rom at Louisiana State University Health Sciences Center-Shreveport, and Drs. Shu Chien and John Shyy at the University of California San Diego for the valuable discussion; Dr. Holly Yin and the High Throughput Screening Core at City of Hope for the technical assistance. Z.B.C., X.L. and K.-C.W. conceived the study, Z.B.C. and K.-C.W. acquired funding, and supervised the experiments and analyses. X.L., Z.B.C., and K.-C.W. designed the experiments and wrote/revised the manuscript. X.L. performed the majority of mouse and cell experiments and the related data analyses. Y.L., P.S., and M.K. designed the AGO1-ASO. Y.L. and X.T. performed the initial *in vitro* and *in vivo* tests of the ASO. X.T. performed the lipid profiling in HFHS diet-fed animals. Y.L. prepared the bulk RNA-seq library. D.Y. prepared Cut&Tag library and analyzed the sequencing data with X.L. and performed immunoblotting and co-IP. A.T. and N.K.M. performed liver EC isolation. D.Y. and N.K.M performed immunofluorescent staining and quantification. R.S.C. performed the SRS imaging under the supervision of L.W., S.T. and N.P. designed and synthesized the LNP for ASO delivery. T.-Y.W. and K.-C.W. designed, synthesized and characterized the MoNP for ASO delivery. All authors have contributed intellectually to the study and reviewed and edited the draft of manuscript.

## Supplemental information

### Supplemental Material and Methods

#### Chow and HFHS diet feeding

Mice were fed a chow diet (D12489B, Research Diets Inc, 16.4% kcal protein, 70.8% kcal carbohydrate, 4.6% kcal fat) or an irradiated HFHS (D12266B, Research Diets Inc, 17% kcal protein, 32% kcal fat, 51% kcal carbohydrate) diet for 16 weeks starting at 8 weeks old.

### EC isolation

Liver EC isolation was performed as described with modifications.^73^ Briefly, mice were humanely euthanized, the vascular system washed with PBS via cardiac perfusion and livers were collected. In a small tube, the liver tissue was chopped into small pieces and blood cells were lysed using ACK lysis buffer. Next, tissue was digested in collagenase type I at 37°C for 60 minutes. ECs were enriched by using mouse anti-CD146 and CD144 antibodies (REAfinity, Miltenyi Biotec, CA, USA). For aortic ECs, aortas were collected after perfused with PBS and then flushing with TRIzol reagent (Thermo Fisher Scientific), as described^71^.

### Isolation of Bone marrow-derived monocytes and PBMC

Bone marrow-derived monocytes were isolated from mouse femurs and tibias using the Monocyte Isolation Kit (BM), mouse (Miltenyi Biotec, Cat# 130-100-629) following published protocol.^74^ Briefly, bone marrow cells were flushed with cold MACS buffer, gently dissociated by pipetting, filtered to remove cell clumps, and washed. Cells were incubated with FcR Blocking Reagent and Monocyte Biotin-Antibody Cocktail, followed by Anti-Biotin MicroBeads. Magnetically labeled non-monocyte cells were depleted using MACS separation columns, and the unlabeled monocyte-enriched flow-through fraction was collected, washed, and used for qPCR analysis.

PBMC was isolated from mouse whole blood by density-gradient centrifugation using Lympholyte-M (Cedarlane, Cat# CL5035).^75^ Briefly, blood was collected into heparinized PBS, diluted with 1× PBS, and layered onto Lympholyte-M, followed by centrifugation at 1,700 rpm for 25 minutes. The PBMC layer was removed, washed twice in 1× PBS, and used for qPCR analysis.

### Plasma biochemical index testing

Total, HDL, and LDL/VLDL-cholesterol were measured in serum using the HDL and LDL/VLDL Quantitation Kit (Sigma-Aldrich #MAK045; #CS0005) following manufacturer’s instructions. Specifically, 15-50 μl of serum was fractionated by adding equal volume of fractionating reagent (2x LDL/VLDL Precipitation Buffer Sigma#MAK045B) and then centrifugation at 12,500g for 10 minutes to yield the HDL supernatant phase and a pellet containing the low-density lipoproteins (LDL and VLDL). These fractions were used to measure cholesterol, and the values were added to obtain the total circulating cholesterol.

ALT was measured in serum using the Liquid ALT kit (Pointe Scientific # A7526-01-1953), and AST was measured in serum using the Aspartate Aminotransferase Colorimetric Activity Assay kit (Cayman #701640). The change in absorbance in both assays was recorded for 40 minutes at 37°C, calculating the activity from the slope from a minimum linear period of 10 minutes.

### Atherosclerotic and liver lesion analysis

After humane euthanasia, mice were perfused with 20 mL 0.01 M PBS through the left ventricle. The total aorta and heart or carotid arteries were harvested and fixed. The *en face* aortic root preparation and quantification of the lesion areas in the whole aorta were performed as described.^76^ For *en face* analysis, the whole aorta was cut open and stained with oil red O (ORO) (Sigma). Lesions in the aortic arch (aortic root to below the left subclavian artery), thoracic aorta (the region between the end of the arch and the last intercostal branch) and abdominal aorta (the region between the end of the thoracic aorta segment and the iliac bifurcation) were assessed. For aortic root analysis, the heart was embedded in OCT, frozen in liquid nitrogen, and cross-sectioned serially at the aortic root level at a 7 μm thickness. For carotid arteries, after the gross images were taken, the LCA was cut into two segments for cross and longitudinal sections.^7^ The cryosections were stained with ORO and counterstained with hematoxylin. Images were taken using Zeiss Observer II microscope, and quantification was determined with ImageJ. Livers from mice were collected, embedded in OCT and sectioned with a cryostat. Slides were fixed in 4% (vol/vol) paraformaldehyde (PFA) for 10 minutes for histological analyses.

### Histology and immunohistochemistry staining

#### Hematoxylin & Eosin staining (H&E)

Tissues were harvested and fixed in 10% neutral buffered formalin. Dehydration, clearing, and paraffinization was performed on a Tissue-Tek VIP Vacuum Infiltration Processor (SAKURA). The samples were embedded in paraffin using a Tissue-Tek TEC Tissue Embedding Station (SAKURA) and sectioned and put on positively charged glass slides. The slides were deparaffinized, rehydrated, and stained with Modified Mayer’s Hematoxylin and Eosin Y Stain (America MasterTech Scientific) on a H&E Auto Stainer (Prisma Plus Auto Stainer, SAKURA) according to standard laboratory procedures.

#### Immunohistochemistry (IHC)

The IHC stains were performed using a Ventana Discovery Ultra IHC automated stainer (Ventana Medical Systems, Roche Diagnostics, Indianapolis, USA). Slides were incubated with endogenous peroxidase activity inhibitor and antigen retrieval solution. F4/80 (Cell Signaling Technology, Cat# 70076, 1:2000), alpha-smooth muscle actin (Abcam, Cat# ab124964, 1:4000), LYVE1 (Cell Signaling Technology, Cat# 98947, 1:200), AGO1 (ThermoFisher, Cat# PA5-50654, 1:200), Luciferase (Abcam, Cat# ab21176, 1:200), CD31 (Epitomics, Cat# AC-0083A, 1:200), CYP2F2 (Santa Cruz, Cat# 374540, 1:100), HAMP (Abcam, Cat# 190775), CYP1A2 (Santa Cruz, Cat#53241) and GS (Sigma, Cat#, G2782) were incubated followed by DISCOVERY anti-rabbit HQ and DISCOVERY anti-HQ-HRP (Ventana). The antibodies were visualized by DISCOVERY ChromoMap DAB Kit (Ventana) and counterstained with hematoxylin and cover slipped. Whole slide images were acquired with a NanoZoomer S360 Digital Slide Scanner (Hamamatsu) and viewed by NDP.view image viewer software.

#### Masson’s trichrome staining

The stains were performed on a BenchMark Special Stains Instrument (Ventana Medical Systems, Roche Diagnostics, Indianapolis, USA) using a Trichrome staining kit (06521908001). The slides were then rinsed, dehydrated, and mounted.

### Stimulated Raman scattering (SRS) microscopy configuration, ratiometric image processing, and data analysis

The SRS microscope setup was as described previously.^77^ Briefly, SRS images were acquired using an 80 μs pixel dwell time that resulted in an acquisition speed of 8.52 s/frame for a 320×320 pixel field of view. The pump beam wavelength was tuned to 791.3 nm for the –CH_3_ channel (2940 cm^-1^), 797.3 nm for the –CH_2_ channel (2845 cm^⁻1^), and 938.3 nm for the phosphate channel (960 cm^-1^). The Stokes beam was fixed at 1031.2 nm.

Liver sections were imaged with 45 mW Pump and 41 mW Stokes power at the sample. Aorta sections were imaged with 45 mW pump and 41 mW Stokes power for the protein channel, and 45 mW pump and 122.5 mW Stokes power for the lipid and phosphate channels at the sample. Image analysis and color coding were performed using ImageJ. Linear unmixing of the protein (–CH_3_) and lipid (–CH_2_) channels was performed in MATLAB (R2024b, MathWorks) using the method as reported.^78^ For ratiometric analysis, a mask image was generated by adjusting the threshold and normalizing non-zero values to one using the CH_3_ image. Unmixed CH_2_ images were then divided by corresponding unmixed CH_3_ images from the same set, and the resulting ratiometric image was multiplied by the mask image to produce the final CH_2_:CH_3_ ratiometric image.

### Cell culture, treatment, and transfection

Cells were kept at 37°C ventilated with 5% CO_2_ and 21% O_2_. HAECs (at passages 5–8) or HLSECs (within passage 3) or HUVECs (passages 5–8) were cultured in complete M199 medium supplemented with 20% FBS (Sigma, M2520) and 1× antibiotics (penicillin–streptomycin, Gibco, 15140122). HepG2 cells were cultured in DMEM with 10% FBS. In some experiments, cells were treated with oxPAPC (40 μg/mL) (Avanti Polar Lipids, Cat# 870604) or vehicle control for 4 hours. In some experiments, cells were pre-transfected with ASOs using Lipofectamine RNAiMax as described. The medium was changed to M199 containing 15% FBS 4-6 hours after transfection.

### Luminex assay

Cultured medium (500 μl in triplicates) was collected and subjected to Luminex assay using a curated human 14-plex panel (NB1097; Bio-techne) at Analytical Pharmacology Core at City of Hope.

### Monocyte adhesion assay

Monocytes were labeled with CellTracker Green CMFDA Dye (Thermo Fisher Scientific) and incubated with a monolayer of HAEC or HUVECs (4 × 10^3^ cells per cm^2^) for 20 minutes. The non-attached monocytes were then washed off with complete EC growth medium. The attached monocyte numbers were imaged using an Echo Revolve microscope under the 10x lens and green fluorescent channel. Average numbers per condition were calculated using a counter on ImageJ in a blinded fashion from five randomly selected fields of technical duplicates.

### Cell fractionation

Cell fractionation was performed as described.^79^ Briefly, 5-10 million cells were collected, washed with 1 mL of cold PBS containing 1 mM EDTA, and centrifuged at room temperature at 500 × g and the cell pellet was collected. A small portion (10%) of the cell pellet was set aside as the input (total fraction). The remaining pellet was suspended in 200 μL of ice-cold lysis buffer (10 mM Tris-HCl, pH 7.5, 0.05% NP-40, 150 mM NaCl) and incubated on ice for 5 minutes. The lysate was gently pipetted over 2.5 volumes of chilled sucrose cushion (24% RNase-free sucrose in lysis buffer) and centrifuged at 15,000 × g for 10 minutes at 4 °C. The supernatant, representing the cytoplasmic fraction, was collected. The pellet was suspended in 50 μL of the RIPA buffer to obtain the nuclear fraction. Protein concentrations were quantified using the BCA method, and SDS-PAGE was subsequently performed. Cytoplasmic and nuclear markers, β-tubulin (CST, cat # 2416) and Lamin B1 (CST, cat # D9V6H), were used for validation.

### RNA extraction, quantitative PCR, and RNA-seq

Total RNA was extracted from tissues and cells using TRIzol reagent (Invitrogen, cat # 15596026). Extraction from tissue had an additional first step of bead homogenization. cDNAs were synthesized using PrimeScript™ RT Master Mix containing both Oligo-dT primer and random hexamer primers. qPCR was performed with Bio-Rad SYBR Green Supermix using the Bio-Rad CFX Connect Real Time system. 36B4 was used as internal control in mouse and β-actin or 18S in human samples. The primers used in this study are summarized in Table S3.

For RNA-seq, 1 µg of total RNA from three biological replicates was used for library preparation with the KAPA mRNA HyperPrep Kit (Roche, cat # KK8581). mRNA was enriched using mRNA capture beads, and libraries were prepared following the manufacturer’s protocol (KR0960-v6.17). The quality of cDNA libraries was assessed using the Agilent 4200 TapeStation System. Libraries were sequenced using the NovaSeq platform (Illumina) with 150-nt paired-end sequencing, achieving a depth of 50∼100 million read pairs.

### Cut&Tag-seq and Chromatin immunoprecipitation (ChIP)

Cells were collected, centrifuged at 600 × g for 5 minutes, and the cell pellets were retained. For each sample, 500,000 cells were used to prepare libraries following the EpiCypher Cut&Tag protocol with CUTANA™ Cut&Tag Kit (EpiCypher, cat # SKU: 14-1102). Briefly, 10 µL of ConA beads per sample were activated using bead activation buffer. The activated beads were incubated with the cell pellets at room temperature for 10 minutes to bind the cells. Subsequently, 0.5 µg of primary antibody was added to each sample and incubated at 4 °C overnight. IgG (EpiCypher, cat # SKU:13-0024) (negative control) and H3K27ac (EpiCypher, cat # SKU: 13-0059) (positive control) antibodies were used. On the second day, pAG-Tn5 was added to each reaction, and tagmentation buffer was used to activate the Tn5 enzyme for chromatin fragmentation. Following tagmentation, CUTANA high-fidelity 2× PCR Master Mix was used to amplify the DNA fragments under the recommended parameters. Targeted chromatin tagmentation was completed following the epicypher protocol. Libraries were amplified with 14 PCR cycles and purified by single-sided 1.3x AMPure bead purification. Finally, cDNA quality was assessed using an Agilent High Sensitivity DNA chip. Libraries were sequenced on the NovaSeq platform (Illumina) with 150-nt paired-end sequencing, achieving a depth of 5–10 million read pairs.

For ChIP, cells were crosslinked with 1% formaldehyde (Sigma Aldrich, F8775) for 15 min at room temperature, quenched with 0.16 M glycine (pH 7.0) for 5 min at 4°C, and washed in cold PBS. Fixed cells were sequentially washed at 4°C for 10 min with ChIP buffer A (10 mM HEPES, pH 7.6, 10 mM EDTA, pH 8.0, 0.5 mM EGTA, pH 8.0, and 0.25% Triton X-100), followed by ChIP buffer B (10 mM HEPES, pH 7.6, 100 mM NaCl, 1 mM EDTA, pH 8.0, 0.5 mM EGTA, pH 8.0, and 0.01% Triton X-100). Cell pellets were resuspended in sonication buffer (15 mM HEPES, pH 7.6, 140 mM NaCl, 1 mM EDTA, pH 8.0, 0.5 mM EGTA, pH 8.0, 0.1% sodium deoxycholate, and 1% Triton X-100) supplemented with Complete Mini EDTA-free protease inhibitors (Roche), 0.5% SDS, and 0.2% N-lauroylsarcosine, at a final concentration of 1∼5 × 10⁷ cells/mL. Chromatin was sheared using a Bioruptor (Diagenode) at high power to generate DNA fragments of 200∼500 bp. The sonicated chromatin was centrifuged at 12,000 rpm for 10 min at 4°C and diluted fivefold to reduce detergent concentrations. Immunoprecipitations were carried out by incubating the sonicated chromatin with a p65 antibody (CST, cat # 6956S, 1∶500) or a rabbit IgG overnight at 4°C. Equal amounts of Protein A and G Dynabeads (Invitrogen), pre-blocked with BSA (1 mg/mL), were added to the antibody-chromatin mixture and incubated for 4h at 4°C. The beads were then subjected to sequential 10-min washes with Wash A (10 mM Tris, pH 8.0, 1 mM EDTA, pH 8.0, 140 mM NaCl, 0.1% SDS, 0.1% sodium deoxycholate, and 1% Triton X-100), Wash B (Wash A adjusted to 500 mM NaCl), Wash C (10 mM Tris, pH 8.0, 1 mM EDTA, pH 8.0, 250 mM LiCl, 0.5% sodium deoxycholate, and 0.5% IGEPAL CA-630), and TE buffer. Beads were resuspended in 100 μL TE, treated with RNase A (20 μg/mL) at 55°C for 30 min, supplemented with 0.75% SDS and 50 mM Tris, and subjected to overnight reverse crosslinking at 65°C. The eluted DNA was then treated with Proteinase K at 55°C for 2 h and purified using phenol-chloroform extraction. Enrichment fold was determined by using the 2^−ΔCq^ method relative to input controls.

### Seq data analysis

RNA-seq data were analyzed as we previously described.^27^ Raw RNA-seq reads were trimmed to remove adaptors using Trimmomatic (v.0.39) with default settings. Processed reads were aligned to the human reference genome (hg38) using STAR (v.2.7.10) with default parameters. Gene expression was quantified as transcripts per million (TPM) using featureCounts from the Subread package (v.2.0.1). Genes with expression levels below 10 reads were excluded. Fold changes were calculated as the ratio of expression levels in treatment versus control samples. Differential expression analysis was conducted using DESeq2. Genes with false discovery rate (FDR) < 0.05 were considered significant. Expression was classified as upregulated for fold changes >= 0.5 and downregulated for fold changes <=-0.5. The Human MSigDB v2024.1.Hs dataset was downloaded from the GSEA database, and genes were ranked by log fold change in descending order. GSEA enrichment analysis was performed using the clusterProfiler R package. Reads were converted to TPM and scaled using z-scores for heatmap plotting.

Cut&Tag data analysis was performed as published.^80^ Raw sequencing reads with low quality and adaptor sequences were trimmed using Trimmomatic (v.0.39) with default parameters. Trimmed reads were aligned to the hg38 reference genome using Bowtie2 with the parameters “--local -- very-sensitive-local --no-mixed --no-discordant”. Duplicate reads were removed using Picard, and the aligned reads were sorted using Samtools. Normalized bigWig files were generated from BAM files using Deeptools with the “–normalizeUsing RPKM” parameter. For peak calling, MACS2 was used with the following parameters: “callpeak --keep-dup all --p 1e-5 --f BAMPE”. Peaks overlapping blacklist regions were removed using Bedtools intersect. Filtered peaks were annotated to the nearest genes from the peak center using the annotatePeaks.pl script from HOMER. Promoter regions were defined as regions within 3 kb of the transcription start site (TSS). Prior to creating bigWig track files, BAM files were converted to BED format using Bedtools bamtobed. Genome-wide visualization of bigWig files and called peaks was performed using the WashU Genome Browser. Heatmap analysis was conducted using the Deeptools package. Transcription factor enrichment and motif analyses were performed with the HOMER v5.0 software package.

### Western blotting, proximity ligation assay (PLA), co-IP, and (co-)IF

#### Western blotting

Tissues or cells were homogenized in RIPA buffer (50 mM Tris-HCl, pH 8.0, 150 mM NaCl, 5 mM EDTA, 1 mM DTT, 1% NP-40, 0.1% SDS, and 1× protease inhibitor cocktail). Proteins were separated by SDS-PAGE and transferred onto PVDF membranes. Membranes were processed according to the ECL western blotting protocol (ThermoFisher). Images were captured using the Amersham Imager 680 (GE Healthcare). The gray values of Western blot bands were quantified using ImageJ. Antibodies included mouse AGO1 monoclonal antibody (2A7) (WAKO, cat # 015-22411), rabbit AGO1 polyclonal antibody (ThermoFisher, cat# PA5-50654), rabbit β-tubulin (CST, cat # 2144), PCSK9 rabbit mAb (ABclonal, cat# A21909), rabbit NF-κB (D14E12) (CST, cat # 8242), rabbit p-AMPKα (Thr172) (40H9) (CST, cat # 2535), rabbit phospho-acetyl-coA carboxylase (Ser79) antibody (CST, cat # 3661), CPT1α rabbit mAb (ABclonal, cat# A22450), Lamin B1 (D9V6H) Rabbit mAb (CST, cat # 13435), and IκBα polyclonal antibody (Proteintech, cat# 10268-1-AP).

#### Proximity Ligation Assay (PLA)

The Duolink® PLA fluorescence protocol was followed. ECs were seeded onto 4-well glass chamber slides (cat. #PEZGS0416, Millipore) and allowed to grow and underwent transfection and treatment. Cells were blocked with Duolink blocking solution for 60 min at 37 °C and then incubated with primary antibodies (AGO1, Invitrogen, cat. #PA5-50654; NF-κB, CST, cat. #6956) diluted in antibody diluent at 4°C overnight. For the negative control, PLA was performed without primary antibodies. Slides were subsequently incubated with pre-heated Duolink PLUS and MINUS PLA probes in a humidified chamber for 1 h at 37°C. Ligation was performed by adding ligase and incubating for 30 min at 37°C, followed by signal amplification with polymerase for 100 min at 37°C. Slides were then washed with wash buffer and prepared for imaging.

#### Co-IP

For co-IP, cells were lysed in RIPA buffer (10 mM Tris-HCl, pH 8.0, 150 mM NaCl, 1 mM EDTA, 1 mM DTT, and 0.1% NP-40) supplemented with protease and phosphatase inhibitors. The lysates were incubated overnight at 4°C with Protein A/G agarose beads and AGO1 antibody (Thermo Fisher, cat # PA5-50654, 1:1000) or p65 antibody (CST, cat # 8242S, 1:500), while rabbit IgG antibody served as a control. After extensive washing with lysis buffer, the immunoprecipitates were eluted and subjected to SDS-PAGE, followed by Western blot analysis.

#### (Co-)IF

For (co-)IF, ECs seeded on coverslips were washed with PBS and fixed in 4% paraformaldehyde at room temperature for 15 minutes. Cells were permeabilized in PBS containing 0.3% Triton-X-100 for 10 minutes, rinsed with PBS, and blocked with 10% goat serum at room temperature for 1 hour. Coverslips were incubated with primary antibodies anti-AGO1 (Thermo Fisher, cat # PA5-50654, 1∶1000) only or also with anti-p65 (CST, cat # 6956S, 1∶1000) diluted in 10% goat serum overnight at 4°C. Cells were washed with PBS and subsequently treated with anti-Rabbit Alexa Fluor™ 594 (Thermo Fisher, cat # A-11012, 1∶1000) or anti-mouse Alexa Fluor™ 488 (Thermo Fisher, cat # A-10680, 1∶1000) secondary antibodies at room temperature for 1 hour. Coverslips were washed and mounted with mounting media containing DAPI. IF images were captured using a Zeiss confocal microscope and a PerkinElmer Operetta CLS High Content Analysis imaging system. Images were analyzed in Fiji (ImageJ) by analyzed by two independent researchers in a blinded manner. Briefly, channels were separated and DAPI-positive nuclei were segmented by automated thresholding, hole filling, watershed separation, and particle analysis. For AGO1/p65 co-localization studies, individual nuclei were defined as regions of interest (ROIs), and Pearson’s correlation coefficients were calculated on a per-nucleus basis using pixel intensities from the AGO1 and p65 channels. For AGO1 nuclear localization analysis, p65-positive cell masks and DAPI-positive nuclear masks were used to define cellular, nuclear, and cytoplasmic compartments, and AGO1 mean fluorescence intensity and integrated density were quantified to calculate nuclear-to-cytoplasmic ratios.

## Supplemental Figures

**Figure S1.**
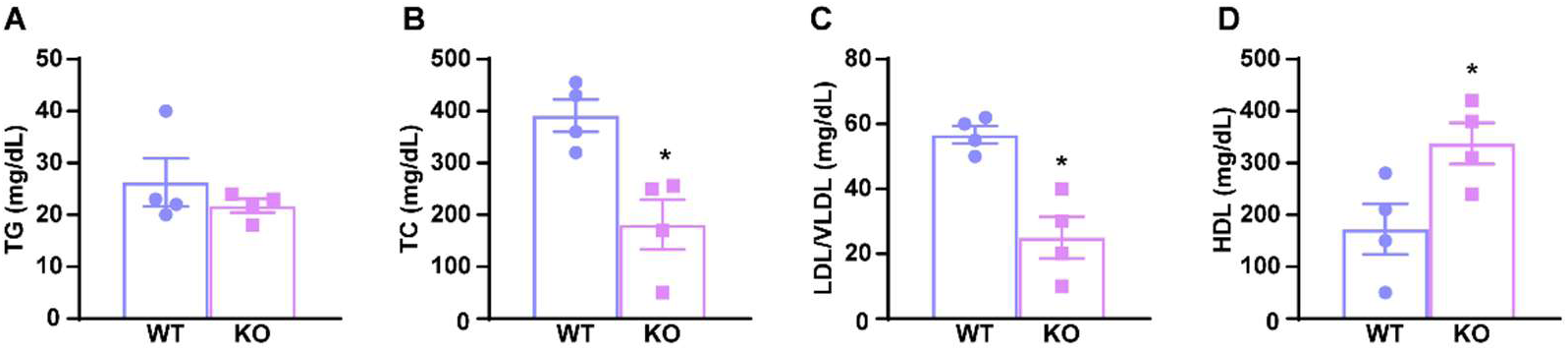
EC-AGO1-KO improves plasma lipid profiles in a diet-induced obesity model. Starting at 8 weeks old, male EC-AGO1-KO mice and WT littermates were fed a HFHS diet for 16 weeks (n=4/group). **A-D**, Plasma triglyceride, total cholesterol, LDL/VLDL, and HDL levels after 4 hours fasting. Data are presented as mean±SEM. *p<0.05 based on Student’s t-test.

**Figure S2.**
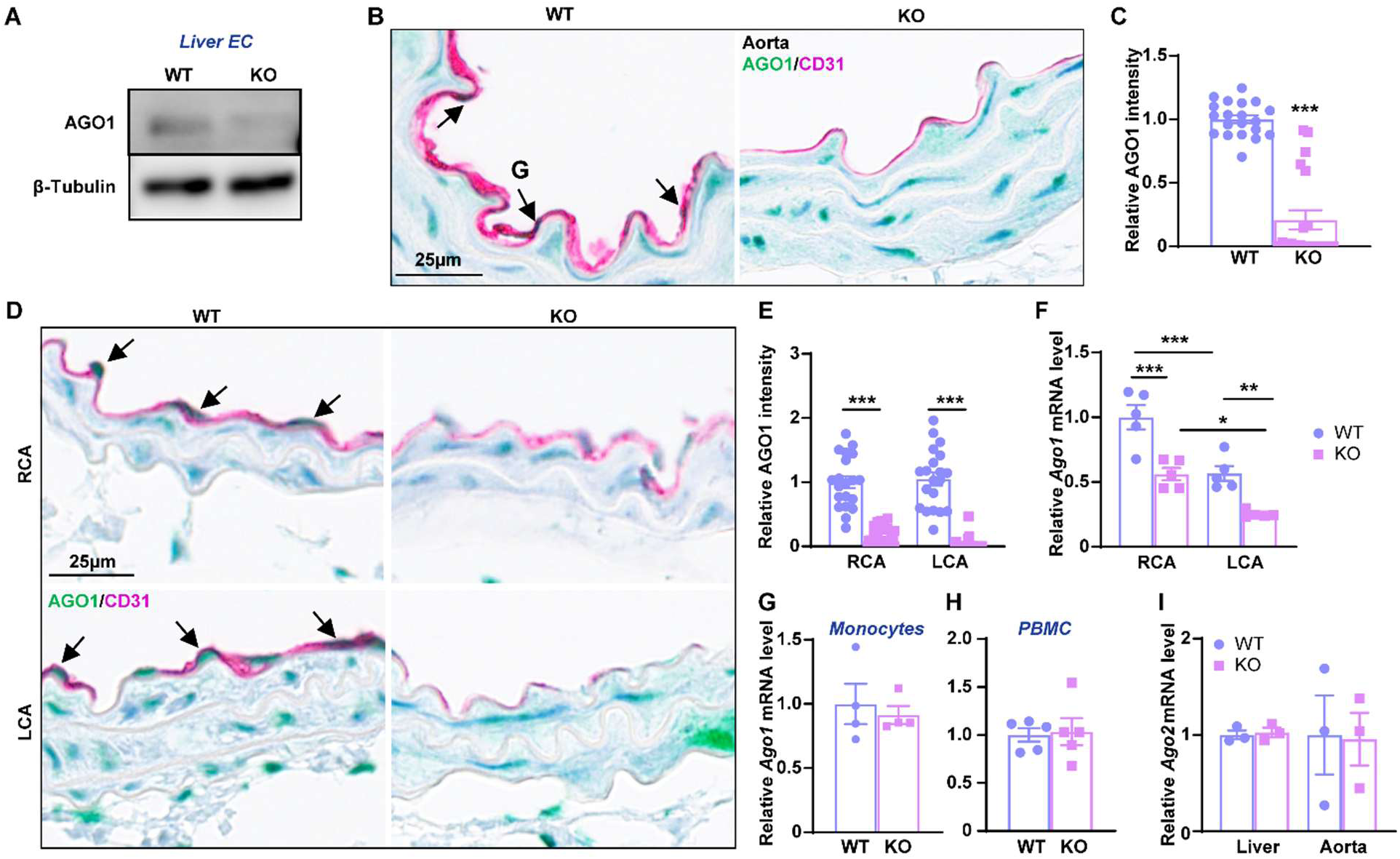
AGO1 expression in liver and aortic EC, monocytes, and PBMCs and *Ago2* expression in ECs from the EC-AGO1-KO vs WT littermates. A, Western blotting for AGO1 protein in liver EC isolated from 12-week-old male mice. **B**, **C**, Representative images of AGO1 (green) and CD31 (pink) dual IHC staining of aorta (in **B**) and quantification of AGO1 in CD31^+^ cells (in **C**) (n=20 cells per group). **D**, **E**, Representative images of AGO1 (green) and CD31 (pink) dual IHC staining of right carotid artery (RCA) and left carotid artery (LCA) (in **D**) and quantification of AGO1 in CD31^+^ cells (in **E**). **F**, qPCR analysis of mRNA levels of *Ago1* in RCA and LCA (in **F**), bone marrow-derived monocytes (in **G**) and peripheral blood mononuclear cells (PBMCs) (in **H**) and *Ago2* in ECs isolated from the liver and aorta (in **I**) from 12-week-old male EC-AGO1-KO and their WT littermates (n=3-5/group). 36B4 was used as an internal control. Data are presented as mean±SEM, *p<0.05, **p<0.01 and ***p<0.001 based on Student’s t-test (**C**, **G**, **H**) and two-way ANOVA (**E**, **F**).

**Figure S3.**
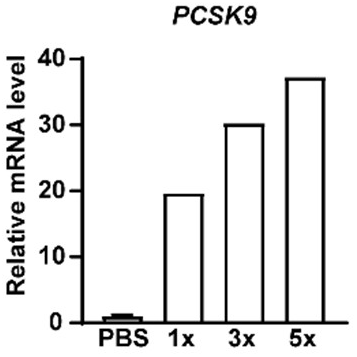
Dose-dependent expression of AAV9-PCSK9. WT mice were injected with PBS or 1×, 3×, and 5×10^11^ viral genome (vg) of AAV9-PCSK9. Livers were collected 1-week later. mRNA levels of PCSK9 were quantified by qPCR with 36B4 as the internal control.

**Figure S4.**
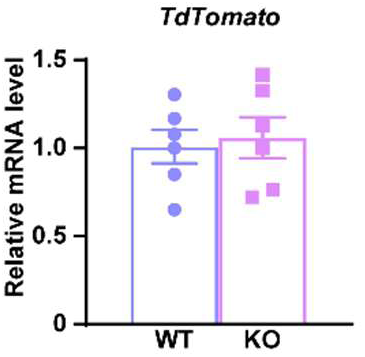
EC-AGO1-KO show no effect on AAV transduction efficiency. Male EC-AGO1-KO mice and WT littermates were injected with the same dose of AAV9-tdTomato through tail-vein and the livers were collected 1-week later. mRNA levels of *Tdtomato* were quantified by qPCR with 36B4 as the internal control (n=6/group). Data are presented as mean±SEM.

**Figure S5.**
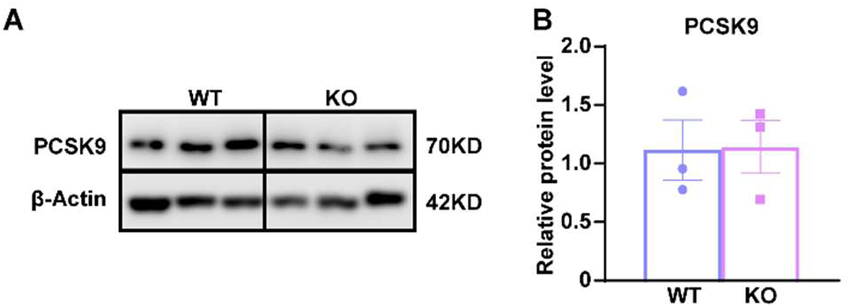
EC-AGO1-KO and WT mice show comparable levels of PCSK9. **A-B**, Liver samples from mice described in Figure 1 underwent Western blotting. Representative image (**A**) and quantification (**B**) of Western blotting for PCSK9. n=3/group. Data are presented as mean±SEM, based on Student’s t-test (**B**).

**Figure S6.**
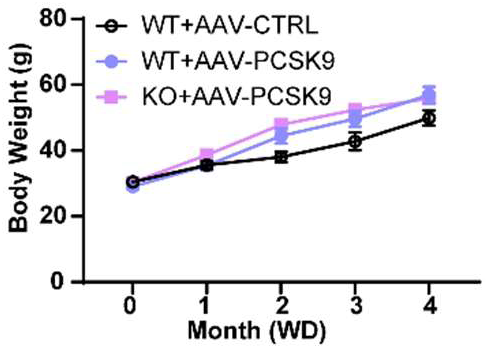
Body weight of EC-AGO1-KO and WT littermates in the AAV-PCSK9 induced AS mice model. Body weight of mice described as in Figure 1 (n=5-7/group). Data are presented as mean±SEM.

**Figure S7.**
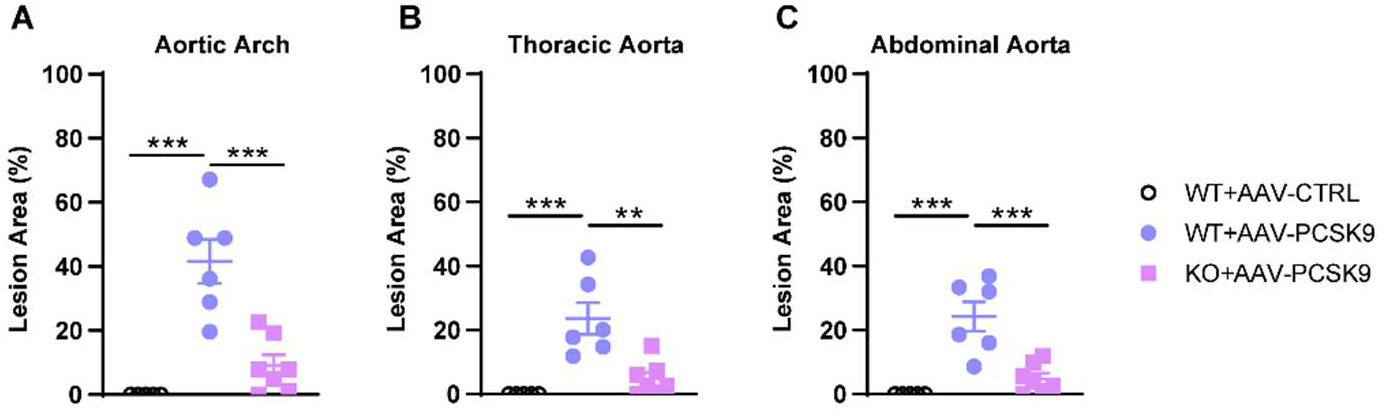
EC-AGO1-KO attenuates atherosclerosis in different segments of aorta in the AAV-PCSK9 and atherogenic diet exposed mice. Mice were the same as in Figure 1. Quantification of plaque areas in different segments of aorta. Data are presented as mean±SEM. **p<0.01, ***p<0.001 between the indicated groups based on one-way ANOVA.

**Figure S8.**
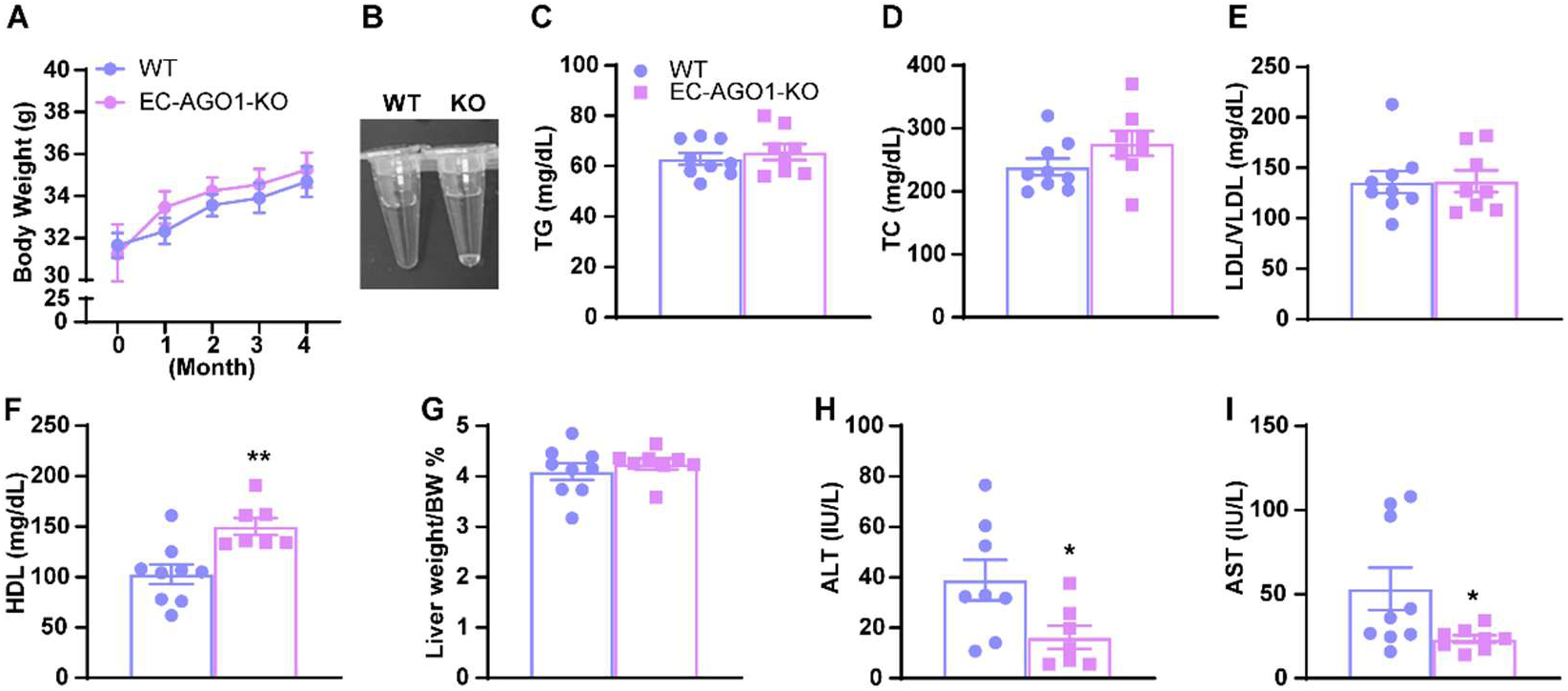
EC-AGO1-KO maintains metabolic homeostasis in AAV-PCSK9-exposed mice. Male EC-AGO1-KO mice and WT littermates were injected with one dose of AAV-PCSK9 and fed chow diet for 16 weeks starting at 12 weeks old (n=9 WT and 8 KO). **A**, Body weight measured monthly from the AAV-PCSK9 administration. **B**, Representative picture of plasma samples from two groups of mice after 4 hours fasting. **C-F**, Plasma triglyceride, total cholesterol, LDL/VLDL and HDL after 4 hours fasting. **G**, Relative liver weight. **H-I**, Plasma ALT and AST levels after 4 hours fasting. Data are presented as mean±SEM. *p<0.05, **p<0.001 based on Student’s t-test.

**Figure S9.**
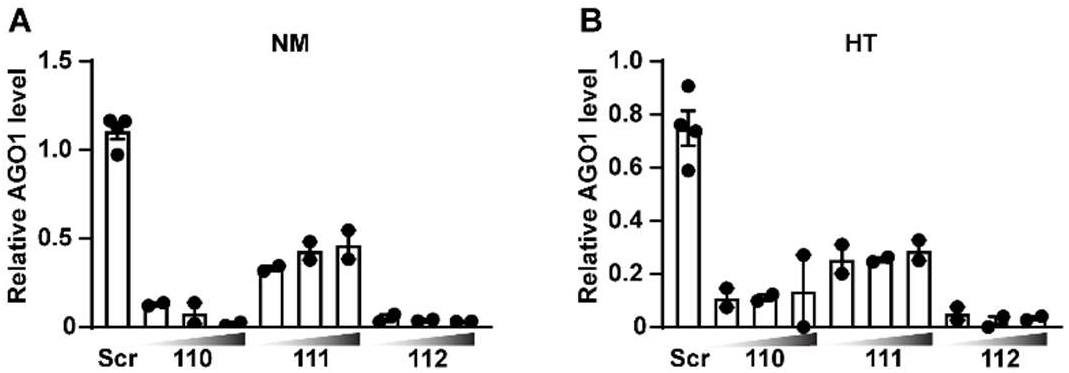
Knockdown (KD) effect of AGO1-ASO in ECs. HUVECs were transfected with a scrambled (Scr) or one of the three AGO1 ASOs (i.e., 110, 111 and 112) at 20, 50 and 100 nM (left to right). *AGO1* mRNA levels were quantified by qPCR, with β-actin as the internal control. Data represents mean±SEM.

**Figure S10.**
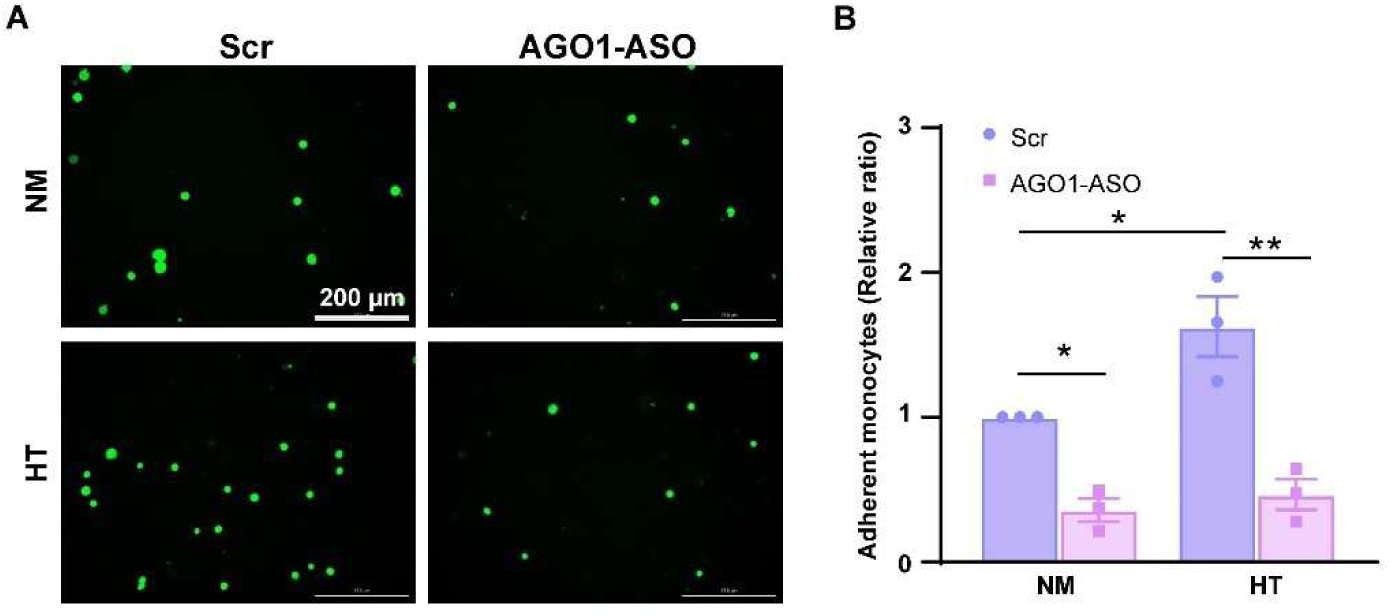
AGO1-KD attenuates monocyte adhesion to HUVECs. HUVECs were transfected with Scr or AGO1-ASO (20 nmol/L) and then treated (HT) for 48 hours for monocyte adhesion assay. Representative images (in **A**) and quantification (in **B**) of monocytes attachment to HUVECs. Scale bar = 200 µm. Data are presented as mean±SEM. *p<0.05, **p<0.01 based on two-way ANOVA.

**Figure S11.**
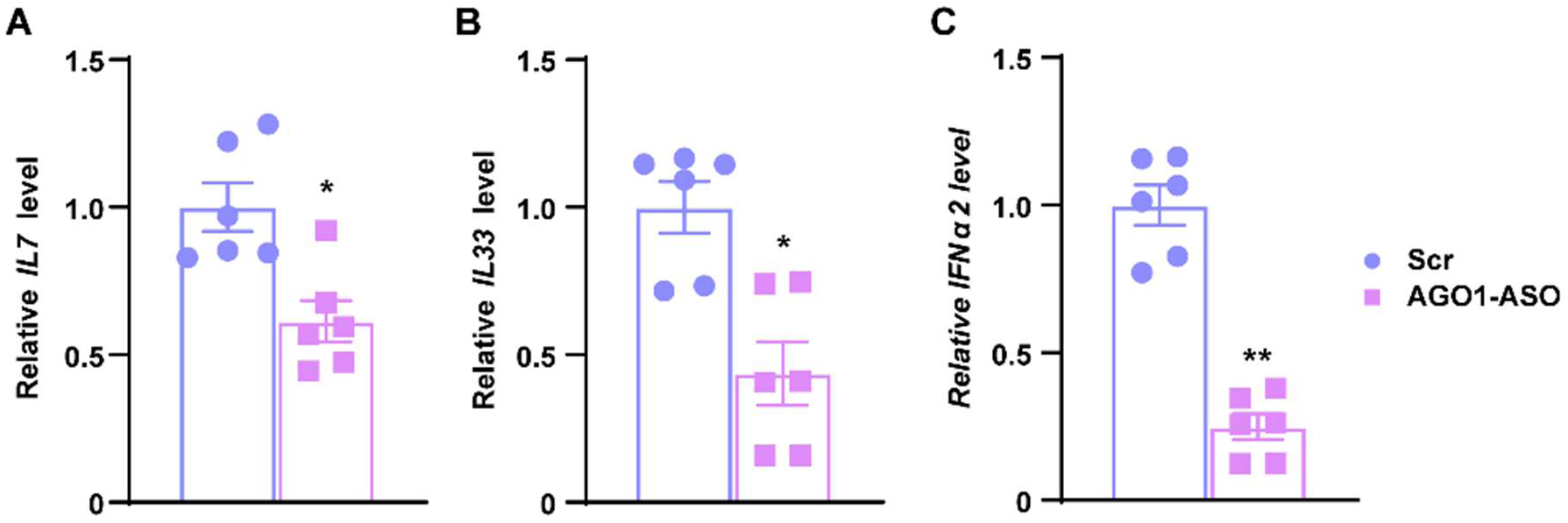
Inhibition of AGO1 in ECs alters gene expression in HAECs. HAECs were transfected with ASO then treated with oxPAPC (40 µg/mL) or a vehicle control for 4 hours. **A**-**C**, The mRNA levels of various genes were quantified by qPCR (n=6). Data are presented as mean±SEM., *p<0.05, **p<0.01 based on Student’s t-test.

**Figure S12.**
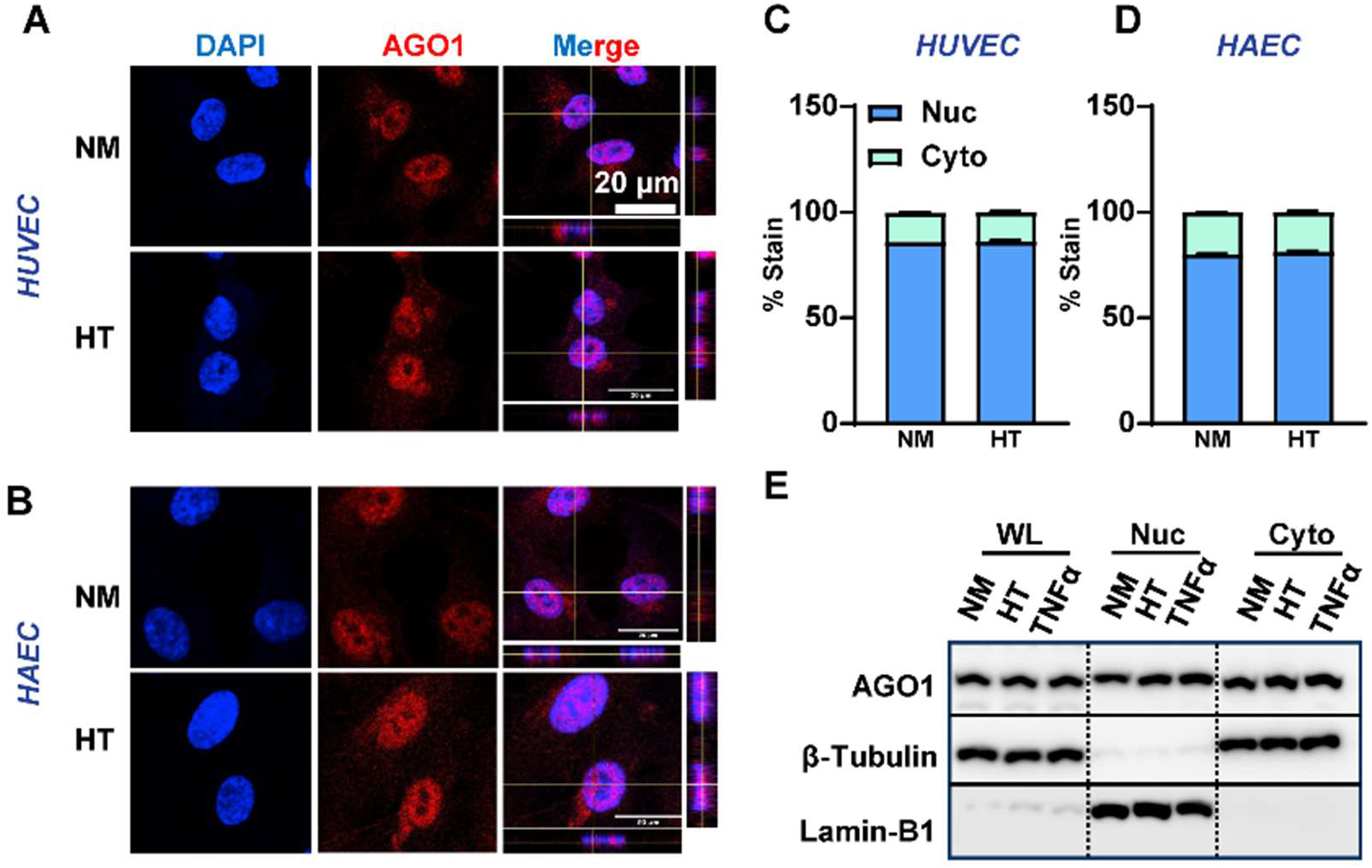
AGO1 shows both cytoplasmic and nuclear location. **A-E**, HUVECs and HAECs were treated by 25 mM mannitol as normal glucose and osmolarity control (NM) or 25 mM D-glucose and 5 ng/mL TNF-α (HT) (**A-D**), or 5 ng/mL TNF-α only (**E**). Confocal imaging of AGO1 immunofluorescent (IF) staining was performed with DAPI counterstain (**A**, **B**) and the subcellular localization was quantified (**C**, **D**). Scale bar=20 µm. **E**, Representative Western blot of AGO1 in subcellular fractions, β-tubulin and Lamin-B1 were used as cytoplasmic and nuclear markers. WL=whole cell lysates, Nuc=nuclear extracts, and Cyto=cytoplasmic extracts. Data are presented as mean±SEM.

**Figure S13.**
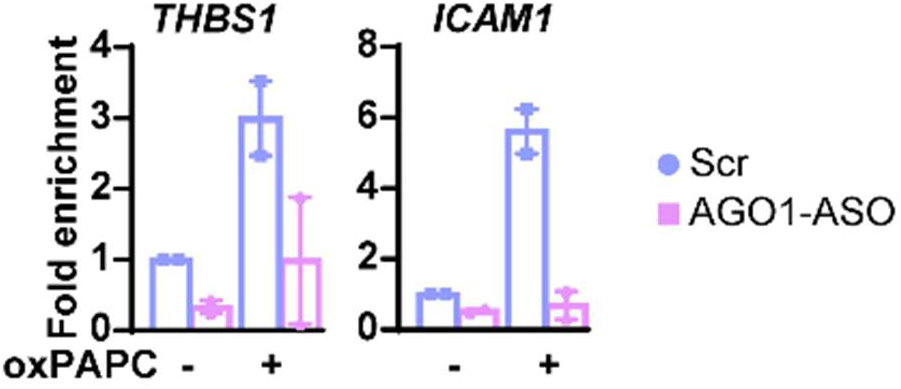
AGO1 binding at the promoter regions of *THBS1* and *ICAM1*. qPCR of AGO1 binding at the promoter regions of *THBS1* and *ICAM1* performed with Cut&Tag samples from Figure 4D**, E**. Data are presented as mean±SEM.

**Figure S14.**
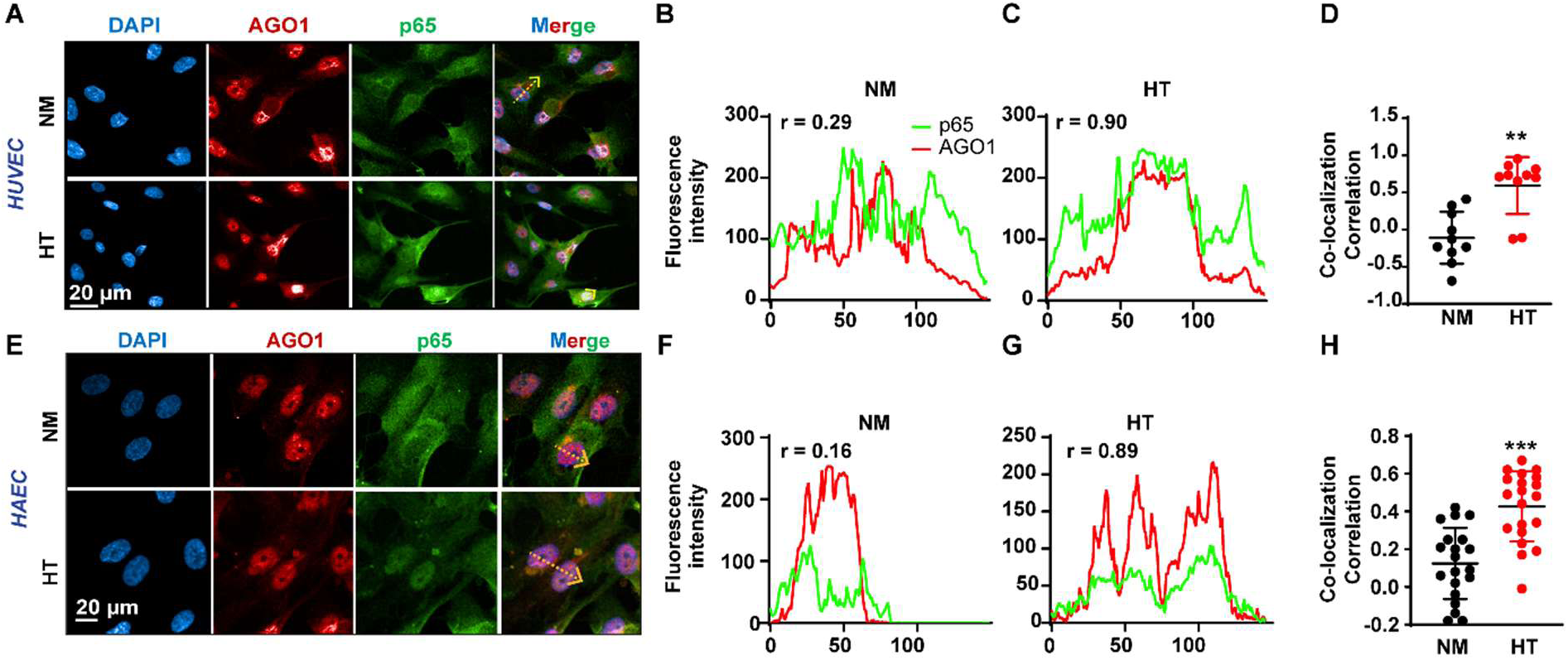
Co-IF of AGO1 and NF-κB in HUVECs and HAECs. **A**, **E** Representative confocal microscopy images of AGO1 (red) and p65 (green) in HUVECs and HAECs treated with HT. The arrow in each merged image indicates the plane for generating line profiles and calculation of Pearson’s correlation coefficient (r) (**B**, **C** and **F**, **G**). **D**, **H**, The co-localization of AGO1 and p65 in the nucleus were quantified in 10 cells (in **D**) and 20 cells (in **H**) per group. Scale bar = 20 µm in **A** and **E**. Data are presented as mean±SEM. **p<0.01 and ***p<0.001 based on Student’s t-test.

**Figure S15.**
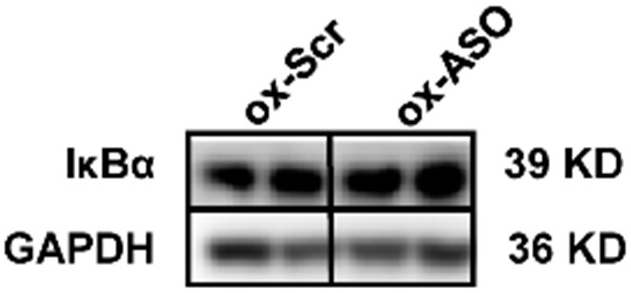
IκBα protein level in HAECs. HAECs were transfected with Scr or AGO1-ASO, (20 nM), before treated with oxPAPC (40 μg/mL) for 4 hours. Representative images of Western blotting of IκBα.

**Figure S16.**
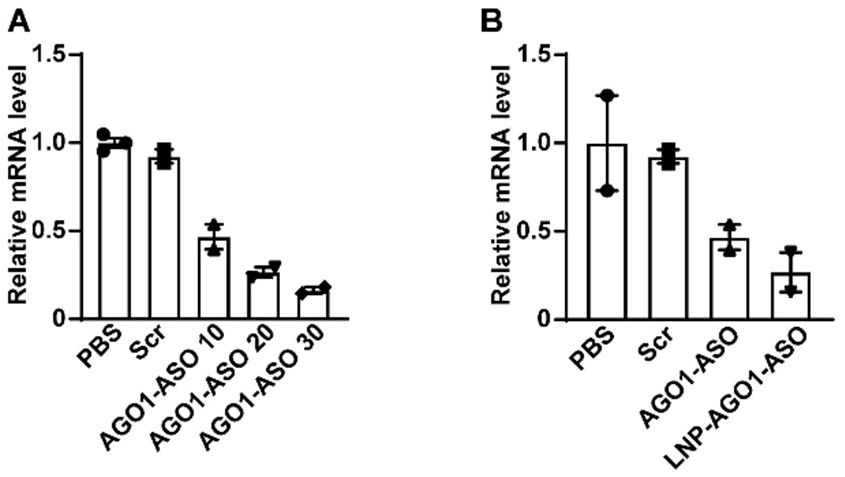
ASO and LNP-ASO AGO1 knockdown efficiency test in mice. **A**, WT mice were injected i.v. with PBS, Scr, or AGO1-ASO at the dose of 10, 20, or 30 mg/kg body weight. A week later, livers were harvested and mRNA levels of *Ago1* were quantified by qPCR in livers after one week. **B**, WT mice were *injected i.v.* with PBS, Scr, AGO1-ASO (10 mg/kg body weight), or LNP-AGO1-ASO (1 mg/kg dose). A week later, livers were harvested and mRNA levels of *Ago1* were quantified by qPCR. n=2-3/group. Data are presented as mean±SEM.

**Figure S17.**
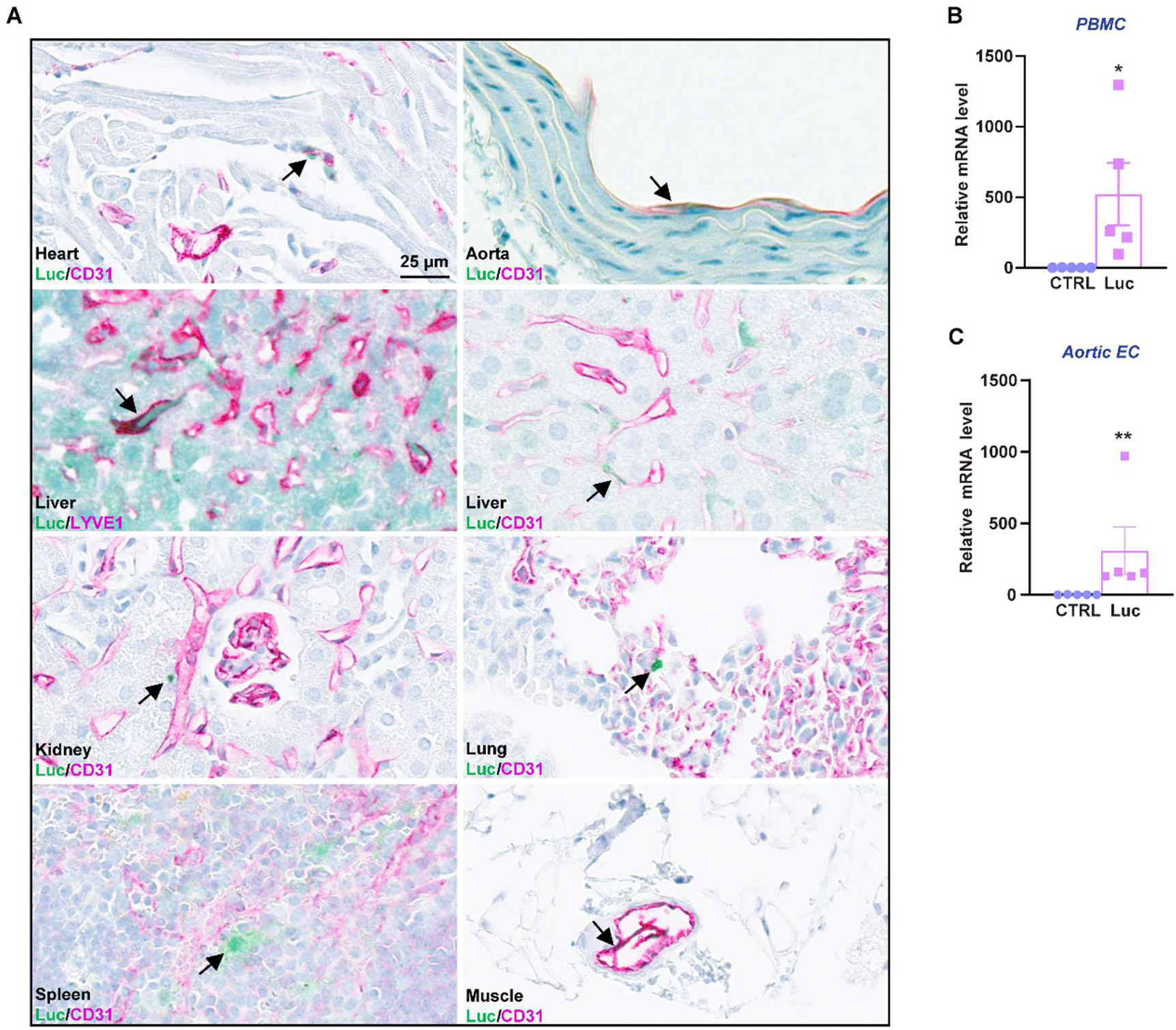
Biodistribution of LNP. LNP containing luciferase mRNA (Luc-mRNA-LNP) were injected through i.v. into WT mice at the dosage of 0.3 mg/kg, control mice were injected PBS (27-week-old, n=5). Plasmas were collected for PBMC isolation, aortas were collected for both aortic EC isolation and aorta IHC staining and other tissues were collected for IHC staining 24 hours later. **A**, Representative images of luciferase labeled LNP (green) and CD31 (pink) dual IHC staining of different tissues. **B**, **C**, qPCR analysis of mRNA levels of *luciferase* in PBMC (in **B**) and aortic EC (in **C**). 36B4 was used as an internal control. Data are presented as mean±SEM, *p<0.05 and **p<0.01 based on Student’s t-test.

**Figure S18.**
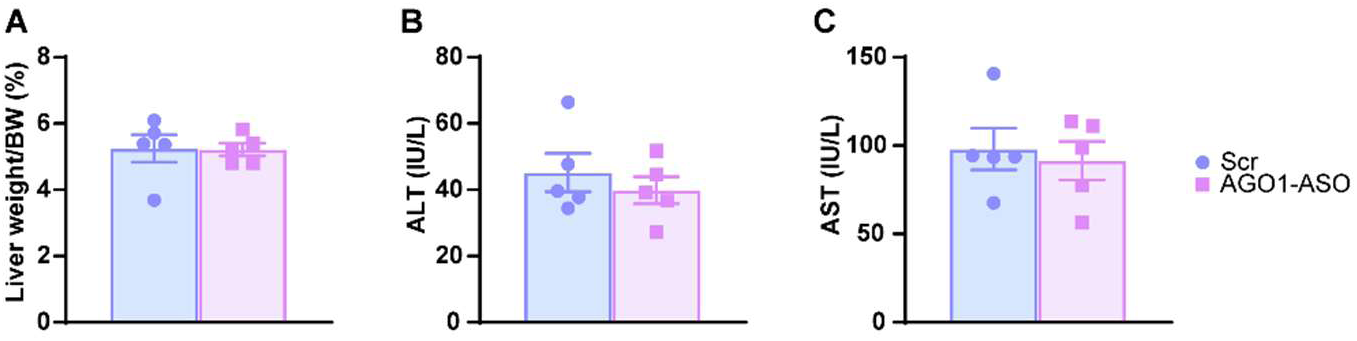
LNP delivery of AGO1 does not show liver toxicity in mice. Mice were treated as in Fig. 5 (n=5 per group). **A**, Relative liver weight. **B, C**, Plasma ALT and AST levels. Data are presented as mean±SEM.

**Figure S19.**
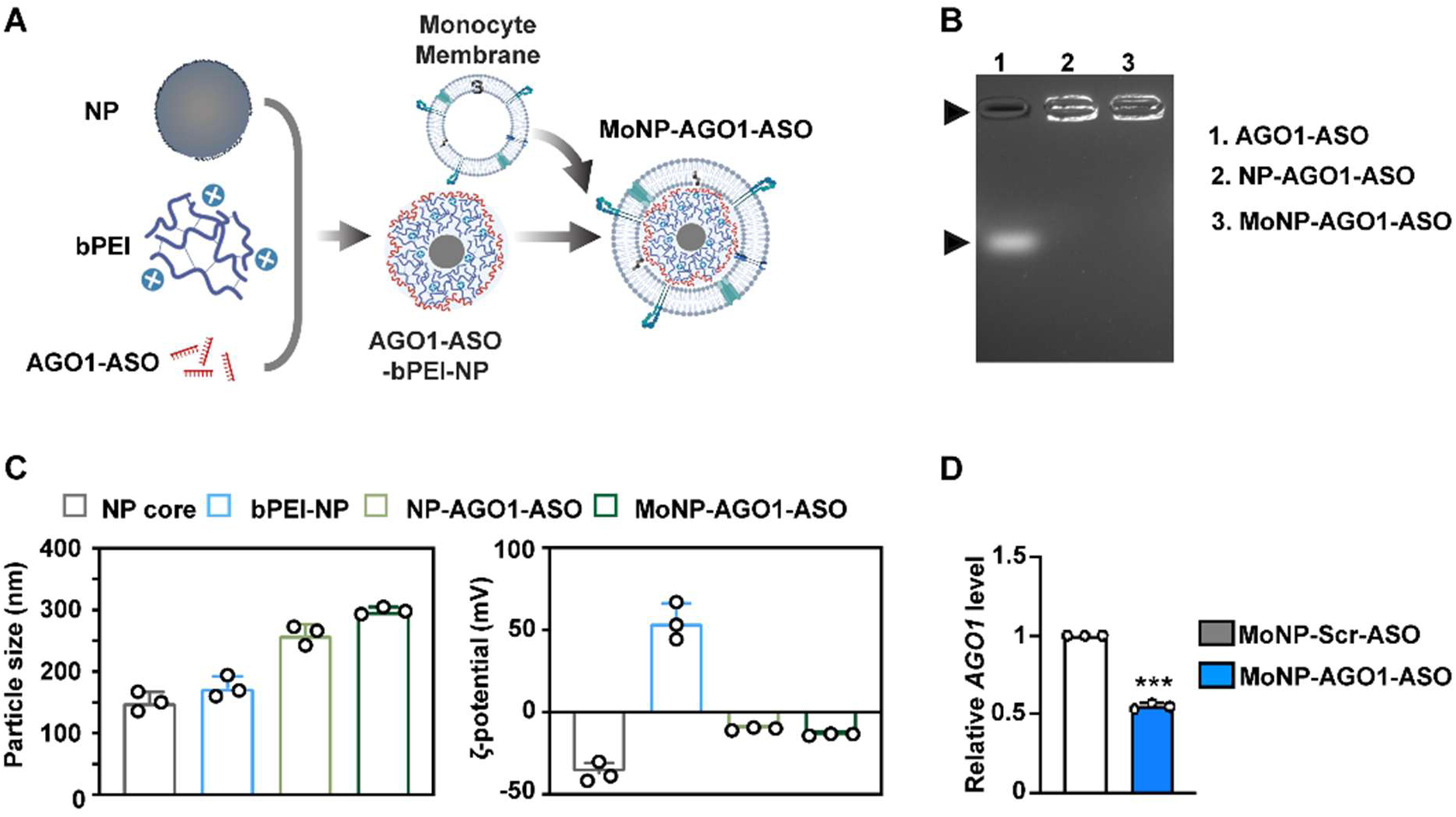
Formulation and characterization of MoNP-AGO1-ASO. **A**, A schematic depicting the formulation process of MoNP-AGO1-ASO. **B**, Gel retardation assay confirming successful incorporation of AGO1-ASO into MoNP. The arrows indicate free AGO1-ASO (lane 1) versus AGO1-ASO conjugated to nanoparticles (lanes 2 and 3). **C**, Dynamic light scattering analysis showing the hydrodynamic size and zeta potential of the resulting MoNP-AGO1-ASO. **D**, HAECs were incubated overnight with MoNP-AGO1-ASO containing 0.1 μg AGO1-ASO or control MoNP, and *AGO1* mRNA level was measured by qPCR analysis. Data are presented as mean±SEM. ***p<0.001 based on Student’s t-test.

**Figure S20.**
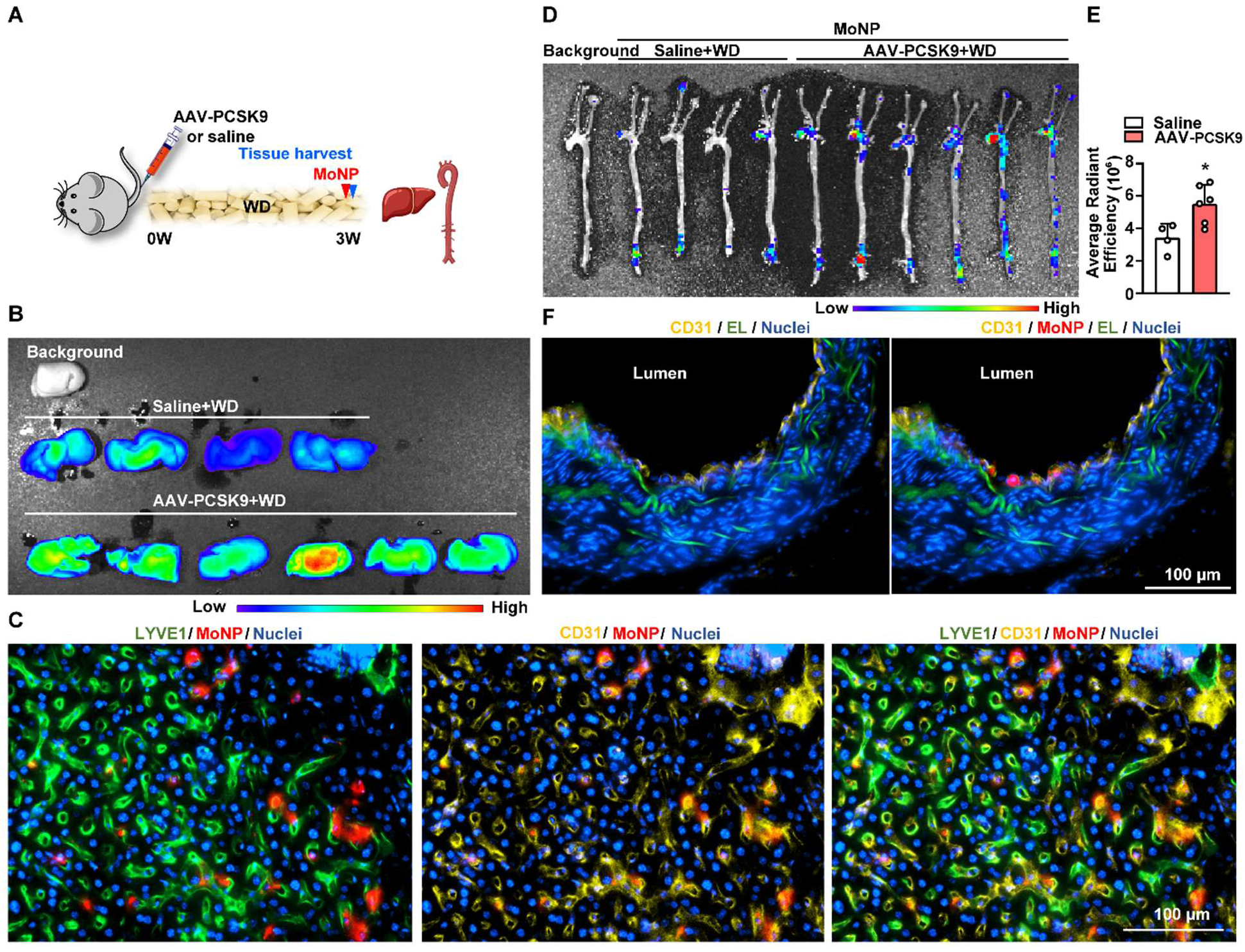
Evaluation of MoNP accumulation in the atherosclerotic mouse aorta and liver. **A**, Experimental design. C57BL/6J mice received AAV-PCSK9 or saline, followed by a WD for 3 weeks. Fluorescently labeled MoNP were intravenously administered 3 hours prior to tissue harvest. **B**, IVIS imaging of mouse livers following MoNP administration. **C,** Representative immunofluorescence images of liver sections. **D**-**E** IVIS imaging of arterial tree and quantification of average radiant efficiency in the image. **F**, Representative immunofluorescence images of aortic arch cross-sections from AAV-PCSK9-treated mice receiving MoNP. In C & F, Red: MoNP, yellow: CD31, green: LYVE1 (C) or elastic lamina (EL) (F), and blue: DAPI. Scale bar = 100 µm. n=4-6/group. Data are presented as mean ± SEM, *p<0.05, based on Student’s t-test (**E**).

**Figure S21.**
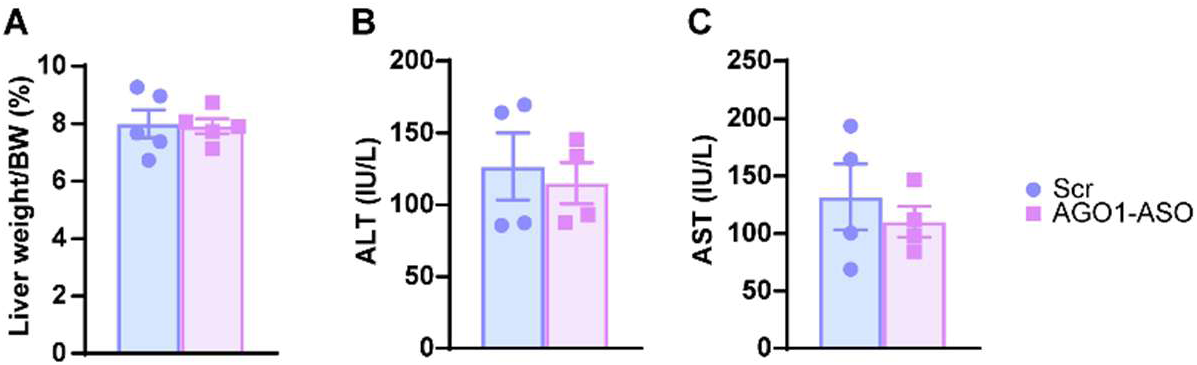
MoNP delivery of AGO1 does not cause liver toxicity in mice. Mice were the same as in Figure 6 (n=5 per group). **A**, Liver weight to body weight ratio. **B**, **C**, ALT and AST levels in plasma. Data are presented as mean±SEM.

**Figure S22.**
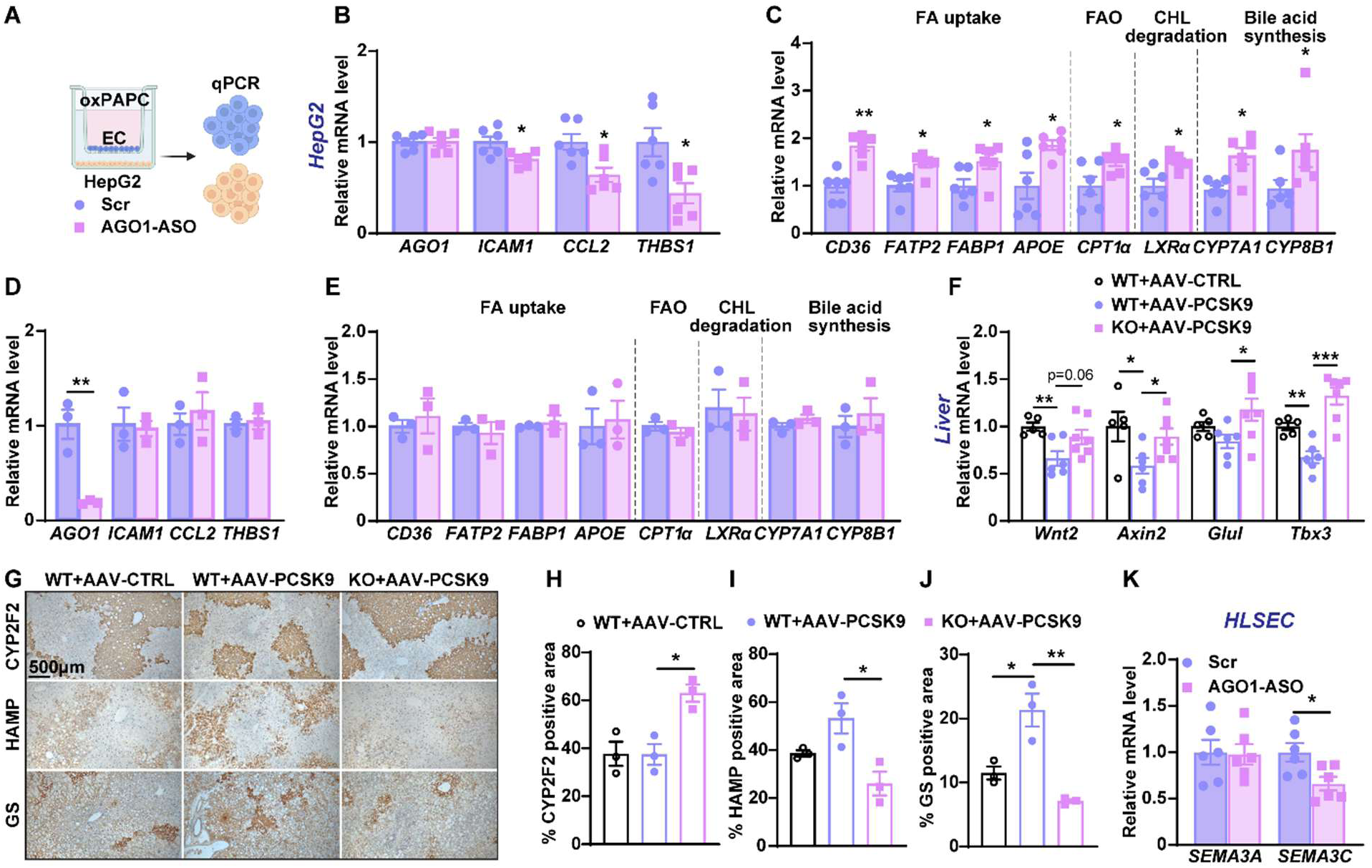
Inhibition of AGO1 in ECs alters gene expression in the hepatocytes and liver. **A**, Experimental design: HLSECs were transfected with ASO then treated with oxPAPC (40 µg/mL) or a vehicle control and co-cultured with HepG2 cells for 4 hours. **B**, **C**, The mRNA levels of various genes in HepG2 cells were quantified by qPCR. **D**, **E**, qPCR of mRNA levels of marker genes associated with inflammation and lipid metabolism in HepG2 cells cultured alone with or without AGO1-KD. **F-J**, Same mice described in Figure 1 were analyzed. **F**, qPCR of mRNA levels of genes involved in Wnt/β-catenin signaling in the liver. **G-J**, Representative images of CYP2F2 (Zone 1), HAMP (Zone 2), and GS (Zone 3) staining of the liver and their quantifications. **K**, The mRNA levels of various genes in co-cultured HLSECs were quantified by qPCR. Data are presented as mean±SEM. *p<0.05, **p<0.01, and ***p<0.001 based on Student’s t-test (**B-E**, **K**) one-way ANOVA (**F**, **H-J**).

**Figure S23.**
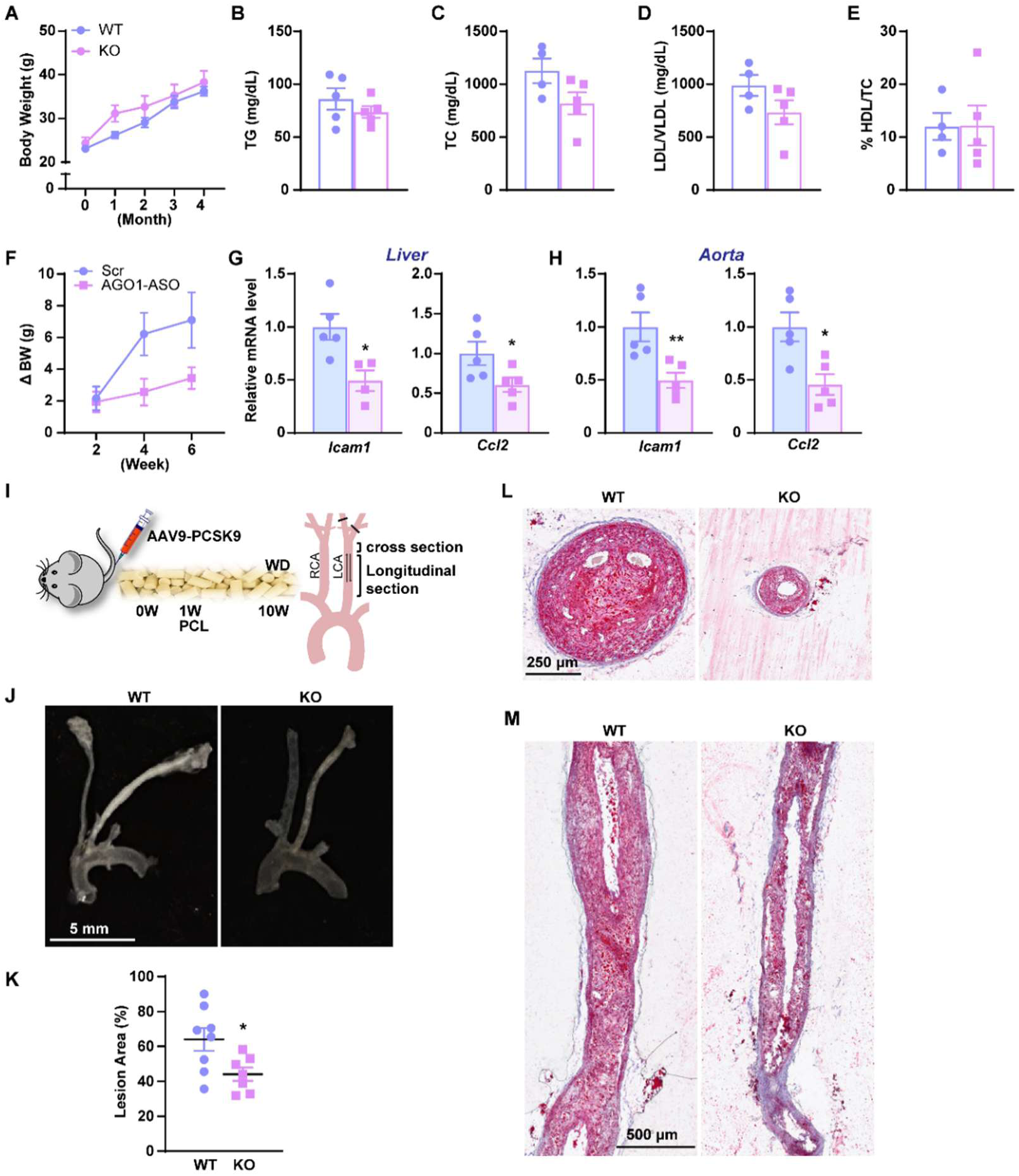
Effect of EC-AGO1-KO and AGO1-ASO in female mice treated by atherogenic regimen. **A-E**, Female mice were under the same treatment as in Figure 1, i.e., injected with AAV9-CTRL/PCSK9 and fed a WD for 16 weeks (n=5 per group). **A**, Body weight. **B-E**, Levels of TG, TC, LDL/VLDL and % HDL/total cholesterol in the plasma after 4 hours fasting. **F-H**, Famle mice were under the same treatment as in Fig. 5, i.e., received AAV9-PCSK9 and fed a WD. Two weeks later, LNP-ASO was i.v. injected at 1 mg/kg body weight weekly for 4 consecutive weeks (n=5 per group). **F**, Body weight. **G**, **H**, qPCR of *Icam1* and *Ccl2* mRNA in livers (**G**) and aortas (**H**). **I-L**, Female C57BL6 WT mice (20-week-old, n=7-8) received AAV9-PCSK9 and were fed a WD. One week later, PCL was performed, LCA and RCA were harvest at 10 weeks. **J**, **K**, Gross LCA image (**J**) and quantification of lesion area (**K**). **L**, **M**, Representative ORO staining images of cross (**L**) and longitudinal sections (**M**) of the LCA (as indicated in **I**). Scale bar = 5 mm in **J**, 250 µm in **L** and 500 µm in **M**. Data represents mean±SEM. *p<0.05, based on Student’s t-test (**A-H** and **K**).

## Supplemental Tables

**Table S1.** Sequences of AGO1 ASOs.

|  |  |
| --- | --- |
| L110 | 5'- +T*+T*+G* A*T*G* G*A*G* A*C*C* T*+T*+A* +A -3' |
| L111 | 5'- +A*+G*+T* C*T*T* C*A*A* T*G*A* T*+C*+T* +C -3' |
| L112 | 5'- +C*+T*+T* G*T*G* T*A*A* G*G*A* A*+T*+G* +T -3' |

\*= phosphothioate +=LNA

**Table S2.**
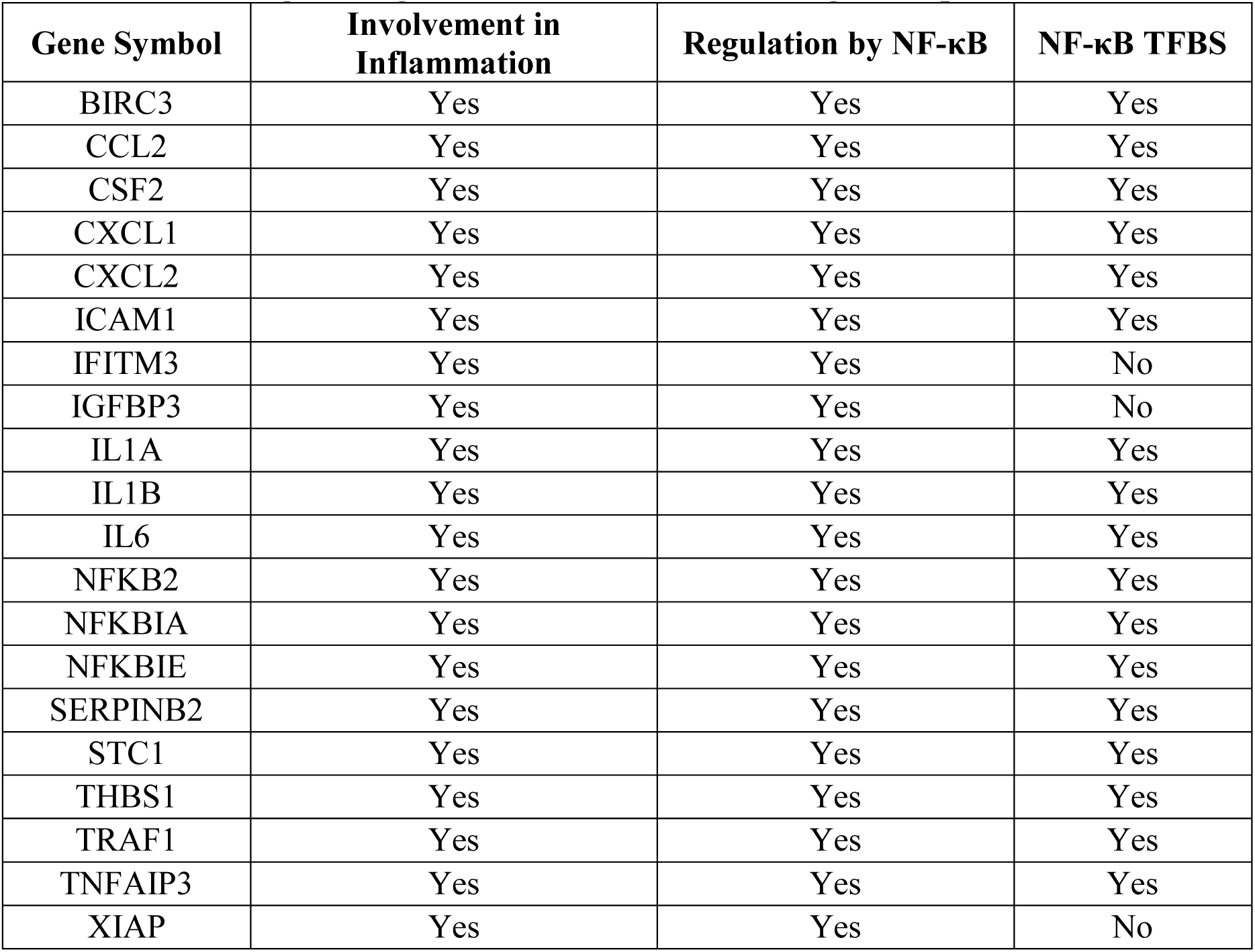
AGO1-regulated genes that show AGO1-binding in the promoters.

**Table S3.**
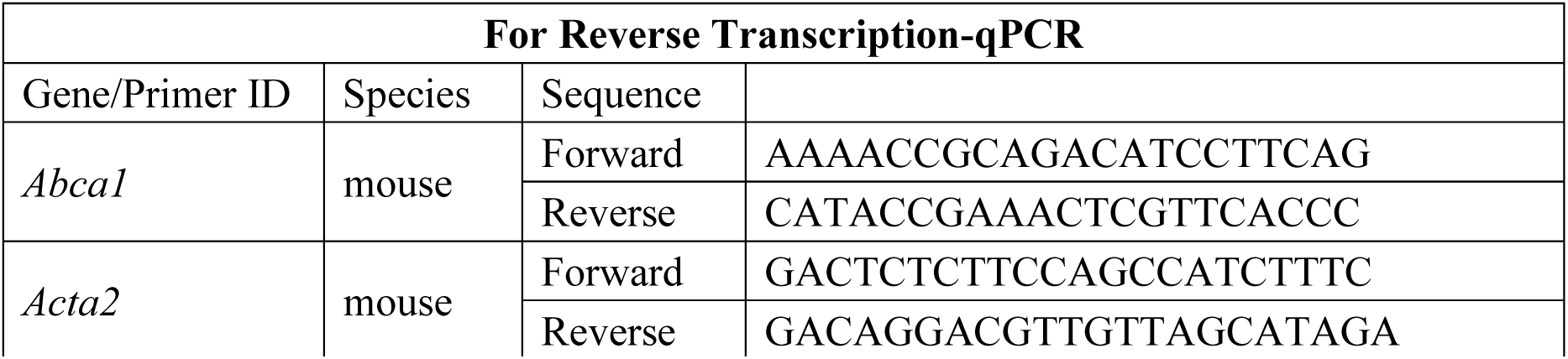
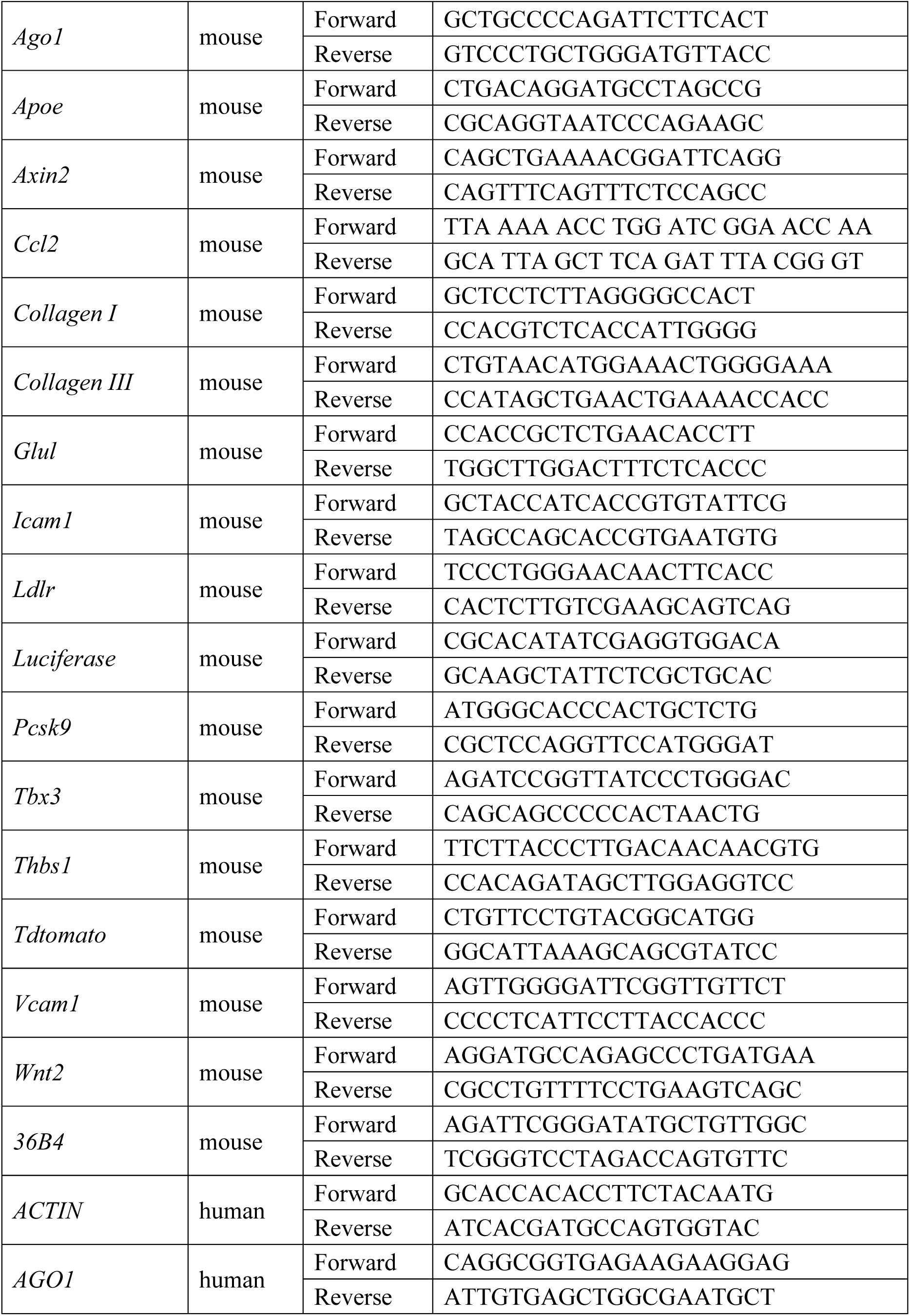
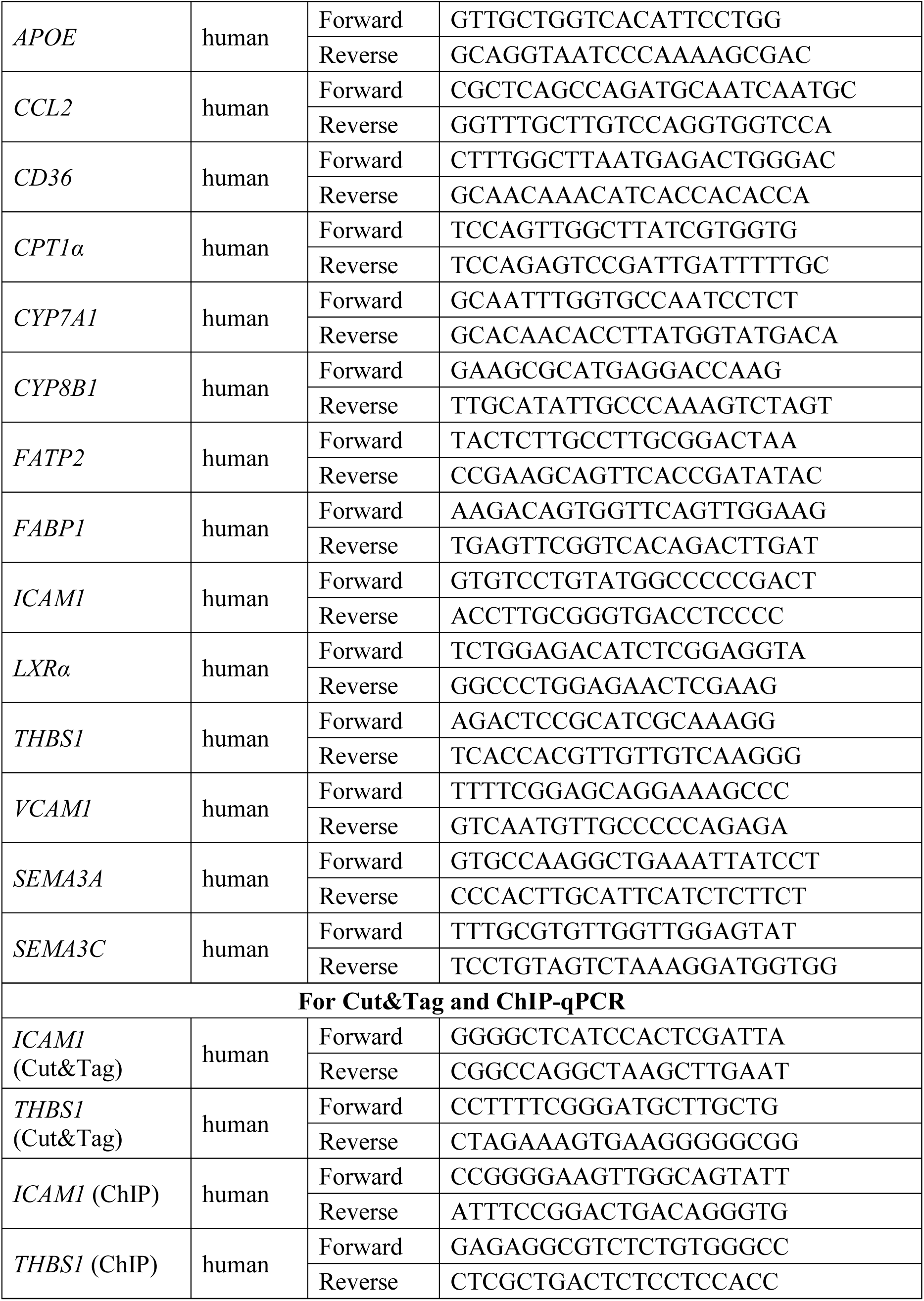
Sequences of primers used for PCR.

